# A starvation-remodeled pre-mRNA structure controls U1 recruitment and nutrient-stress adaptation in yeast

**DOI:** 10.64898/2026.08.06.743327

**Authors:** Jasmine Tsang, Julie Parenteau, Federico Fuchs Wightman, Kristina Sungeun Song, Michelle S. Scott, Silvi Rouskin, Sherif Abou Elela

## Abstract

Recognition of 5′ splice sites by the U1 small nuclear ribonucleoprotein commits pre-mRNAs to splicing, yet splice-site complementarity alone cannot predict productive U1 engagement. Whether dynamic pre-mRNA structure regulates this early spliceosome assembly step remains unclear. Here, we identify a 5′UTR–intron base-pairing interaction positioned near the 5′ splice site that acts as an inducible structural gate for U1 engagement. In budding yeast, this element is enriched among introns required for adaptation to nutrient depletion, and *in vivo* DMS-MaPseq shows that starvation remodels its structure. Structure-guided disruption of pairing impairs adaptation, whereas compensatory mutations restoring pairing without restoring sequence rescue the phenotype, establishing RNA fold as the critical determinant. U1 association decreases when the gate is disrupted and recovers when pairing is restored, and increased Nam8 levels can compensate for gate disruption by stabilizing U1 engagement under stress. Thus, dynamic pre-mRNA folding gates U1 recognition, revealing how transcript architecture converts physiological state into selective splice-site choice.

## INTRODUCTION

Recognition of the 5′ splice site (5′SS) by the U1 small nuclear ribonucleoprotein (snRNP) is the committed step of spliceosome assembly and a key determinant of which pre-mRNAs are spliced and how^1–3^. Although this step is often modeled as base-pairing between the 5′ splice site and U1 snRNA, complementarity alone is a poor predictor of productive U1 association: recognition is a regulated, multi-step process shaped by transcript context and U1-associated factors rather than by duplex strength alone^4–6^. As a result, identical or near-identical 5′ splice-site sequences can be engaged very differently across genes and cellular conditions^7–10^. How information beyond the short splice-site sequence is encoded and read out to control U1 recognition remains a central, unresolved question^4,5^.

This problem is important because U1 recognition is not simply a constitutive housekeeping event. Across eukaryotes, splice-site use changes with development, stress, metabolic state, and disease-associated perturbations of spliceosomal components^11–13^. These observations imply that early spliceosome assembly can be tuned by cellular physiology, yet the molecular features that transmit physiological state to individual splice sites remain incompletely understood. In particular, it remains unclear how local transcript architecture contributes to selective U1 association at native splice sites under changing environmental conditions.

Budding yeast provides a powerful system to address this question. Although *Saccharomyces cerevisiae* has far fewer introns than metazoans, its introns are concentrated in genes with important roles in growth, RNA metabolism, translation, meiosis, and stress adaptation^12,14–17^. The control of meiosis-specific transcripts provides a clear precedent: their splicing is regulated at the stage of 5′ splice-site recognition, showing that U1 engagement, rather than catalysis, can be the decisive regulatory step for a physiological program^18–22^. Yeast splicing is also mediated by spliceosome components that are broadly conserved, including U1 snRNP and its associated RNA-binding factors^23–29^. This combination of a compact intron repertoire, a precedent for U1-level regulation, and conserved machinery makes yeast especially suited to dissecting the rules that govern splice-site recognition *in vivo*.

Nutrient depletion exposes a striking form of spliceosome selectivity in yeast. Under nutrient-rich conditions, highly expressed ribosomal protein transcripts dominate the splicing landscape and occupy a large fraction of spliceosomal capacity^12,30,31^. During starvation, introns, and not simply host-gene expression or protein output, are necessary to confer starvation tolerance and maintain cell function^12,32^. Starvation also changes U1 association and splicing output, redistributing U1 engagement toward a distinct set of intron-containing transcripts that are poorly engaged during normal growth but become important for adaptation to nutrient limitation^12,32^. This redistribution cannot be explained solely by reduced competition from abundant ribosomal protein transcripts, which, even when transcription is repressed, remain more abundant than most other intron-containing RNAs. Instead, the data argue that specific features of starvation-responsive introns promote selective U1 recognition^12,32^. The nature of these features, and how they are regulated by nutrient state, remains unknown.

RNA structure is an attractive candidate for such regulation because it can integrate sequence context, transcript folding, and cellular condition into the local accessibility of regulatory elements. Structured regions can occlude or expose splice sites, recruit RNA-binding proteins, and influence spliceosome assembly in both yeast and metazoans^33–35^. In humans, the choice between competing 5′ splice sites can determine protein output and disease outcome, as illustrated by therapeutic modulation of 5′ splice site recognition in spinal muscular atrophy, where small molecules act in part by stabilizing U1–5′ splice-site complexes^36,37^. More broadly, RNA folding around alternative exons and splice sites has been linked to exon inclusion, splice-site choice, and disease-relevant splicing outcomes in human cells^38–40^. These studies establish RNA structure as an important modulator of splicing, but they do not explain whether physiological stress can remodel a defined pre-mRNA structure near a native 5′ splice site to control U1 recognition.

Here, we use nutrient depletion in budding yeast to ask whether physiological state is encoded in local pre-mRNA structure to control U1 recognition. We identify an inducible 5′UTR–intron base-pairing element near the 5′ splice site that gates U1 engagement, is remodeled by starvation in living cells, and is required for adaptation. These findings reveal that dynamic pre-mRNA folding can convert physiological state into selective splice-site recognition.

## RESULTS

### A 5′UTR–intron structural gate marks introns required for cell tolerance to starvation

During nutrient depletion, yeast cells redistribute U1 snRNP engagement toward a subset of starvation-responsive introns (UpSIs: up-spliced introns during starvation) that support tolerance to starvation, while splicing of many ribosomal-protein and other starvation-dispensable introns (DownSIs: down spliced introns during starvation) is reduced^12^. This intron-dependent fitness requires the 5′ untranslated region (UTR). For at least one intron, MMS2, function depends on a predicted 5′UTR–intron base-pairing interaction near the 5′SS: mutations that unfold this structure abolish function, whereas compensatory mutations that refold it restore function^32^. Whether such a 5′UTR–intron structure is a general, position-defined feature that distinguishes UpSIs from DownSIs, rather than an idiosyncrasy of individual introns, remained unknown.

To address this, we predicted RNA secondary structures spanning the 5′UTR, CDS, and intron for 57 UpSI and 32 DownSI transcripts. Both intron classes formed predicted 5′UTR–CDS–intron contacts, but UpSIs were more frequently folded back toward the 5′ splice site, whereas DownSIs showed more heterogeneous and distal contacts (Figure S1A and S1B). To quantify this difference, we mapped contact frequency across each class. Arc diagrams identified a shared 5′UTR–intron contact cluster near the 5′SS in 74% of UpSIs. By contrast, DownSIs lacked a comparable 5′SS-proximal cluster and instead showed weaker, more dispersed contacts between the 5′UTR and the middle or distal intron (Figure 1A). This positional difference was associated with transcript architecture: UpSIs had longer 5′UTRs and shorter introns than DownSIs, whereas coding-sequence length was similar between groups (Figure S2A and S2B). Consistent with this organization, UpSIs contained a higher fraction of intron nucleotides predicted to pair with the cognate 5′UTR (Figure 1B and S2C). The excess UpSI pairing was concentrated 16–55 nucleotides downstream of the 5′ splice site, defining a 5′SS-proximal region that we refer to as the 5′UTR– intron gate (Figure 1C). This signature was conserved across *Saccharomyces* species: UpSIs showed higher 5′UTR–intron base-pairing scores than DownSIs across nine species, particularly in the gate region adjacent to the 5′SS (Figure 1C). Representative structures illustrate this class difference: the UpSI *SPO1* forms extensive 5′UTR–intron contacts that fold back near the 5′SS, whereas the DownSI *RPL18B* shows more distal contacts and a comparatively accessible 5′SS-proximal region (Figure 1D and 1E). Thus, introns that support starvation tolerance are characterized by a conserved 5′UTR–intron base-pairing gate positioned just downstream of the 5′SS, folding back toward, but not generally encompassing, the splice site, which appears accessible in representative structures. This 5′SS-proximal architecture nominates local pre-mRNA structure as a key determinant of selective 5′ splice-site recognition.

**Figure 1.**
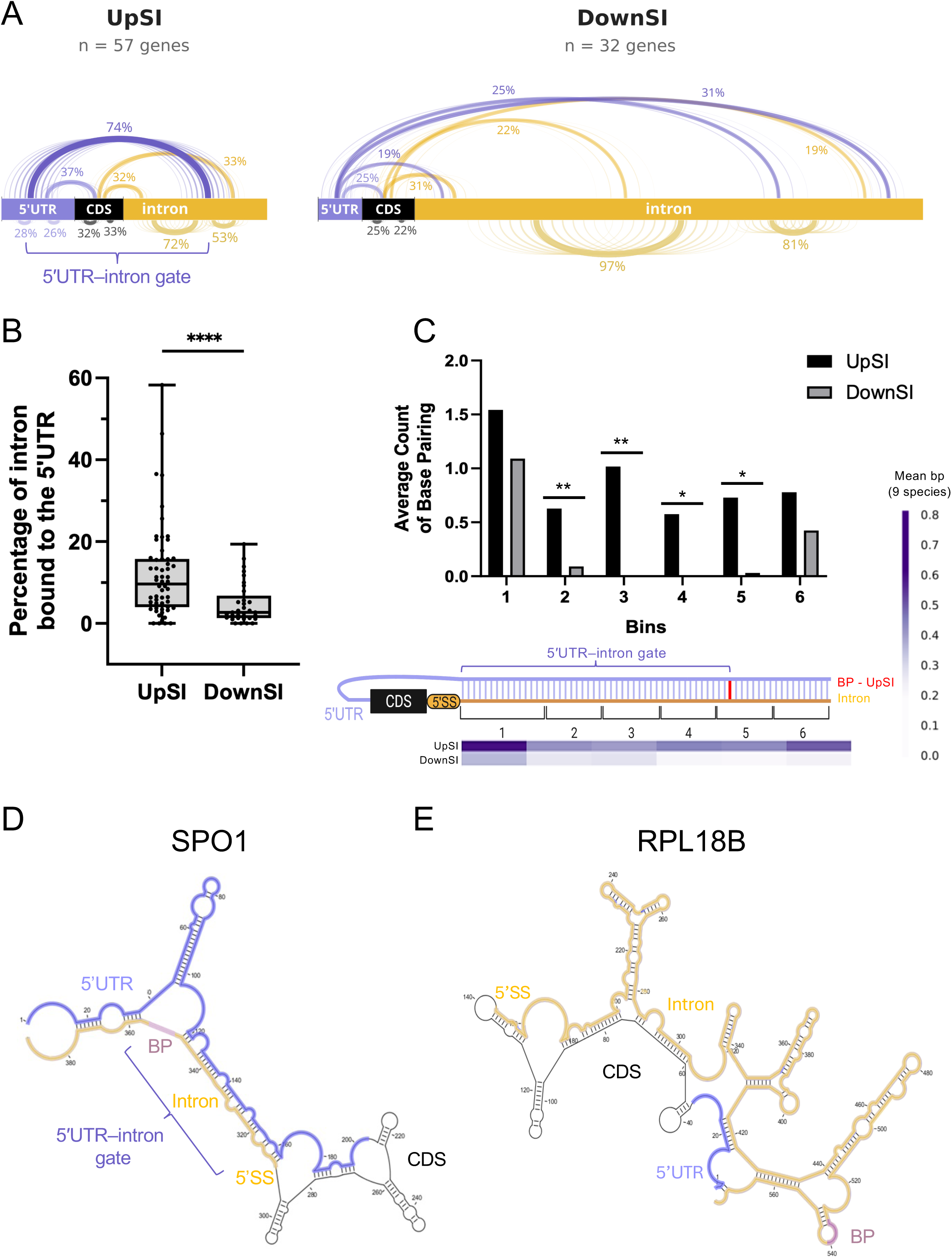
A conserved 5′UTR–intron pairing signature near the 5′ splice site marks starvation-induced introns. **(A)** Arc diagrams of predicted 5′UTR–intron base-pairing contacts in starvation-induced introns (UpSIs; n = 57 genes) and starvation-dispensable introns (DownSIs; n = 32 genes). Region bars indicate the 5′UTR (blue), CDS (black), and intron (orange), scaled to median feature lengths (UpSI: 48, 32, and 94 nt; DownSI: 29, 34, and 334 nt, respectively). Arcs denote contact clusters present in ≥15% of genes; arc thickness and labels indicate the fraction of genes in each cluster, and arc spread reflects positional variability. Only stems of ≥3 consecutive base pairs with pairing probability ≥0.5 were analyzed. **(B)** Percentage of intron nucleotides predicted to base-pair with the cognate 5′UTR in UpSIs and DownSIs. Boxes indicate the median and interquartile range, whiskers indicate the full range, and points represent individual introns. Welch t-test was used as statistical test where ****p<0.0001. **(C)** Average number of 5′UTR–intron base pairs by distance from the 5′SS in *S. cerevisiae*. Bin 1 spans the first 15 nt downstream of the 5′SS, and bins 2–6 each span the next 10 nt. Bars show mean base-pair counts for UpSIs (black) and DownSIs (gray). Significance was determined using a two-sided t-test assuming unequal variances where *p<0.05 and **p<0.01. The heatmap shows mean 5′UTR–intron base-pair counts per gene across nine Saccharomyces species: *S. paradoxus*, *S. kudriavzevii*, *S. mikatae*, *S. cerevisiae*, *S. castellii*, *S. barnettii*, *S. dairenensis*, *S. unisporus*, and *S. servazzii*. The bracketed schematic defines the 5′SS-proximal 5′UTR–intron “gate” region relative to the 5′UTR, CDS, 5′SS, and intron. The red line represents the position of the branch point in UpSI introns. **(D)** Predicted minimum-free-energy secondary structure of the SPO1 UpSI 5′UTR–CDS–intron region. Extensive 5′UTR–intron contacts fold back near the 5′SS, forming a predicted 5′UTR– intron gate. The 5′UTR (blue), CDS (gray), intron (yellow) with specified 5′SS, and branchpoint (BP; purple) are indicated. **(E)** Predicted minimum-free-energy secondary structure of the *RPL18B* DownSI 5′UTR–CDS– intron region. The 5′UTR and intron show more distal contacts, and the 5′SS region remains comparatively accessible. Colors are as in (D). Structures in (D) and (E) were predicted using CLC Main Workbench with default parameters.

### Starvation remodels the UpSI-specific structural gate *in vivo*

Across the analyzed transcript set, predicted UpSI structures had higher, less negative minimum free energy than DownSI structures, indicating that UpSI 5′UTR–intron folds are less thermodynamically stable than DownSI folds (Figure S2D). This lower predicted stability raised the possibility that the UpSI gate could be conformationally responsive, but minimum free energy reflects a predicted ground state and cannot establish RNA folding *in vivo*, where pre-mRNA structure is shaped by RNA-binding proteins, helicases, transcription, and growth state. To determine whether the predicted UpSI gate forms and changes *in vivo*, we probed RNA structure by DMS-MaPseq, in which dimethyl sulfate (DMS) modifies unpaired, solvent-accessible nucleotides and these modifications are read out as mutations by sequencing. Log-phase and 48 h stationary-phase cells were DMS-treated or mock-treated, and modified positions were mapped by reverse transcription, target amplification, and sequencing across the 5′UTR–CDS–intron region of each transcript (Figure 2A). Sequencing libraries showed appropriate per-read mutation burden, uniform amplicon coverage, expected DMS reactivity patterns, and strong signal-to-noise across the analyzed transcripts, supporting downstream structure-guided analysis (Figure S3A–S3D). In *SPO1*, DMS-guided folding revealed nutrient-state-dependent formation of the gate: the log-phase structure showed weak or absent 5′SS-proximal 5′UTR–intron pairing, whereas the stationary-phase structure gained a contact within the gate (Figure 2B). This transition coincided with a local drop in rolling Pearson correlation between log-phase and stationary-phase DMS reactivity profiles, beyond the replicate noise, over the gate region, indicating condition-dependent remodeling rather than preservation of a static fold (Figure 2B). By contrast, the DownSI *RPL18B* showed limited starvation-dependent remodeling near the 5′SS, although it gained an additional intron–intron contact outside the gate-proximal region (Figure 2C). Together, these data show that the 5′UTR–intron gate is remodeled by starvation.

**Figure 2.**
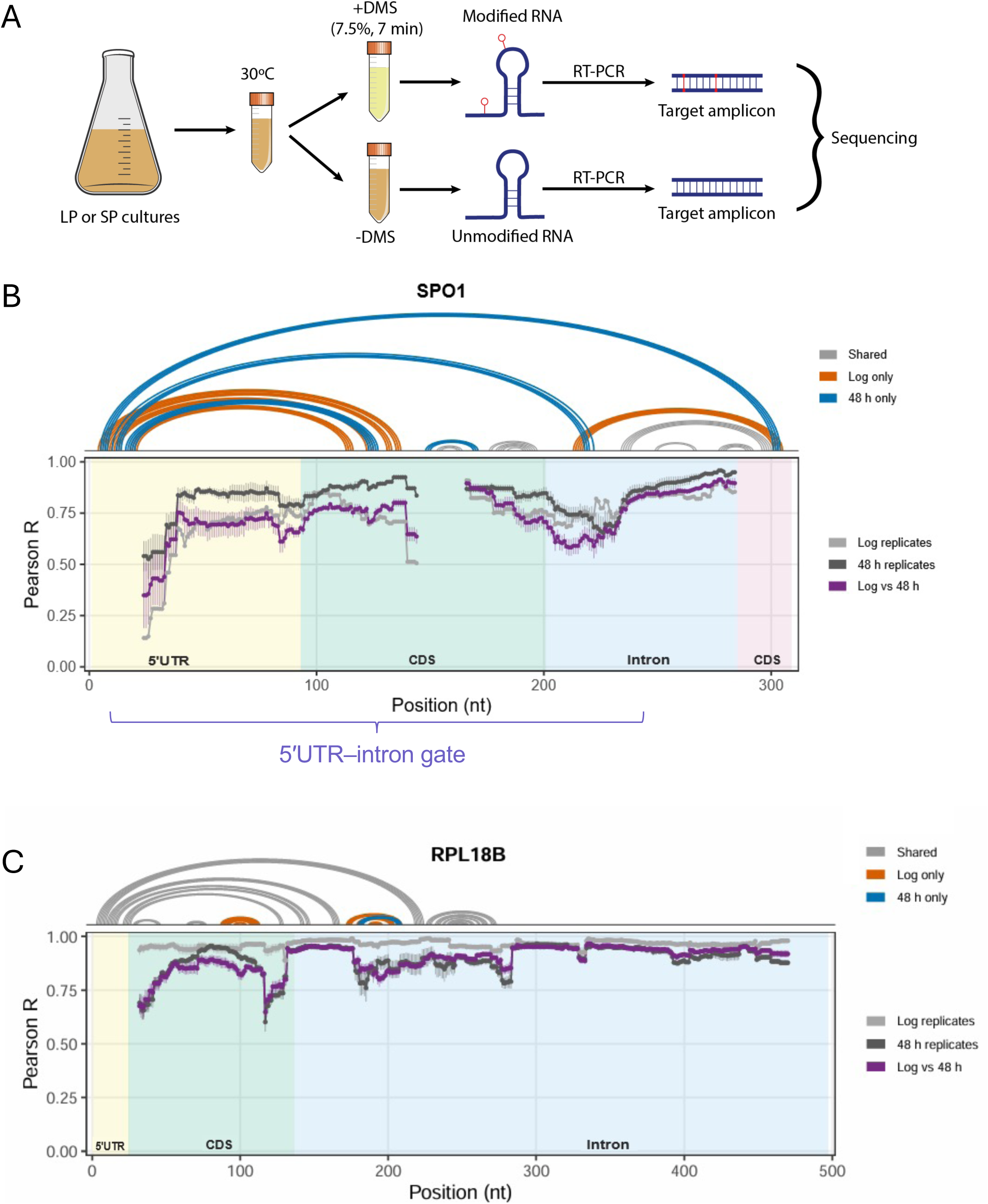
Starvation remodels the *SPO1* 5′UTR–intron gate in cells. **(A)** In-cell dimethyl sulfate mutational profiling with sequencing (DMS-MaPseq) workflow. Log-phase (LP) and 48 h stationary-phase (SP) yeast cultures were treated with DMS or mock treated, followed by RNA extraction, reverse transcription, target amplification, and sequencing to quantify nucleotide-level DMS reactivity. **(B)** In-cell structural remodeling of the SPO1 UpSI 5′UTR–CDS–intron region between log phase and 48 h stationary phase. Top, arc plots show base-pairing interactions detected in log phase only (orange), 48 h stationary phase only (blue), or both conditions (gray). Bottom, rolling Pearson correlation analysis of DMS reactivity profiles using a 45-nt window. Thin traces show replicate comparisons, and bold traces show log-phase replicate correlation (light gray), 48 h stationary-phase replicate correlation (dark gray), and log phase versus 48 h stationary-phase correlation (purple). Shaded regions indicate transcript features, and the bracket marks the 5′UTR–intron gate. **(C)** In-cell structural analysis of the *RPL18B* DownSI 5′UTR–CDS–intron region, displayed as in (B). Compared with SPO1, *RPL18B* shows limited condition-specific remodeling in the 5′SS-proximal 5′UTR–intron region.

### Gate base pairing enables U1 engagement and starvation fitness

To test whether the *SPO1* gate is functionally required, we engineered structure-guided mutations that separate RNA folding from primary sequence. The *spo1-unfold* mutant disrupts the predicted 5′UTR–intron pairing, whereas compensatory substitutions in *SPO1-refold* restore base pairing without restoring the wild-type sequence (Figure 3A). This design allowed us to ask whether the gate acts through its RNA structure rather than through the specific nucleotides that form it. We first measured growth in amino acid-depleted medium containing either 2% or 0.1% dextrose. Under low-dextrose starvation, spo1-unfold phenocopied the intron deletion strain (*spo1Δi*), whereas *SPO1-refold* restored growth toward the wild-type trajectory (Figure 3B). The same mutants grew comparably in 2% dextrose, indicating that gate function is specifically required under the nutrient-limited condition in which the *SPO1* intron supports growth (Figure 3B). Because compensatory refolding rescued the growth defect without restoring the wild-type sequence, starvation fitness depends on the base-paired state of the gate. We next asked whether this growth effect reflected altered U1 snRNP association with *SPO1* RNA. Wild-type and mutant yeast cells expressing an epitope-tagged version of a component of snRNP U1 (TAP-Snu71) were grown in minimal media, crosslinked, and then immunoprecipitated. TAP-Snu71 immunoprecipitation was validated in wild-type and *SPO1* gate-mutant backgrounds by western blot detection of TAP-Snu71 enrichment (Figure S4A). The immunoprecipitated RNA was quantified by RT-qPCR and it showed that starvation increased U1 association with wild-type *SPO1* RNA, but this induction was lost in *spo1-unfold* and restored in *SPO1-refold* (Figure 3C). This effect was not caused by differences in U1 snRNP recovery, because recovery of snR19, the U1 snRNA, remained comparable across strains and conditions (Figure S4D). Thus, U1 association follows the folding state of the *SPO1* gate rather than the wild-type gate sequence. The gate also controlled starvation-associated *SPO1* RNA processing. In wild-type cells, starvation shifted *SPO1* RNA output toward mature mRNA; this shift was impaired in *spo1-unfold* and restored in *SPO1-refold* (Figure 3D). Together, these data establish gate base pairing as the structural signal that converts nutrient depletion into productive U1 engagement and *SPO1* RNA maturation, revealing local pre-mRNA folding as a causal mechanism for selective splice-site recognition during starvation.

**Figure 3.**
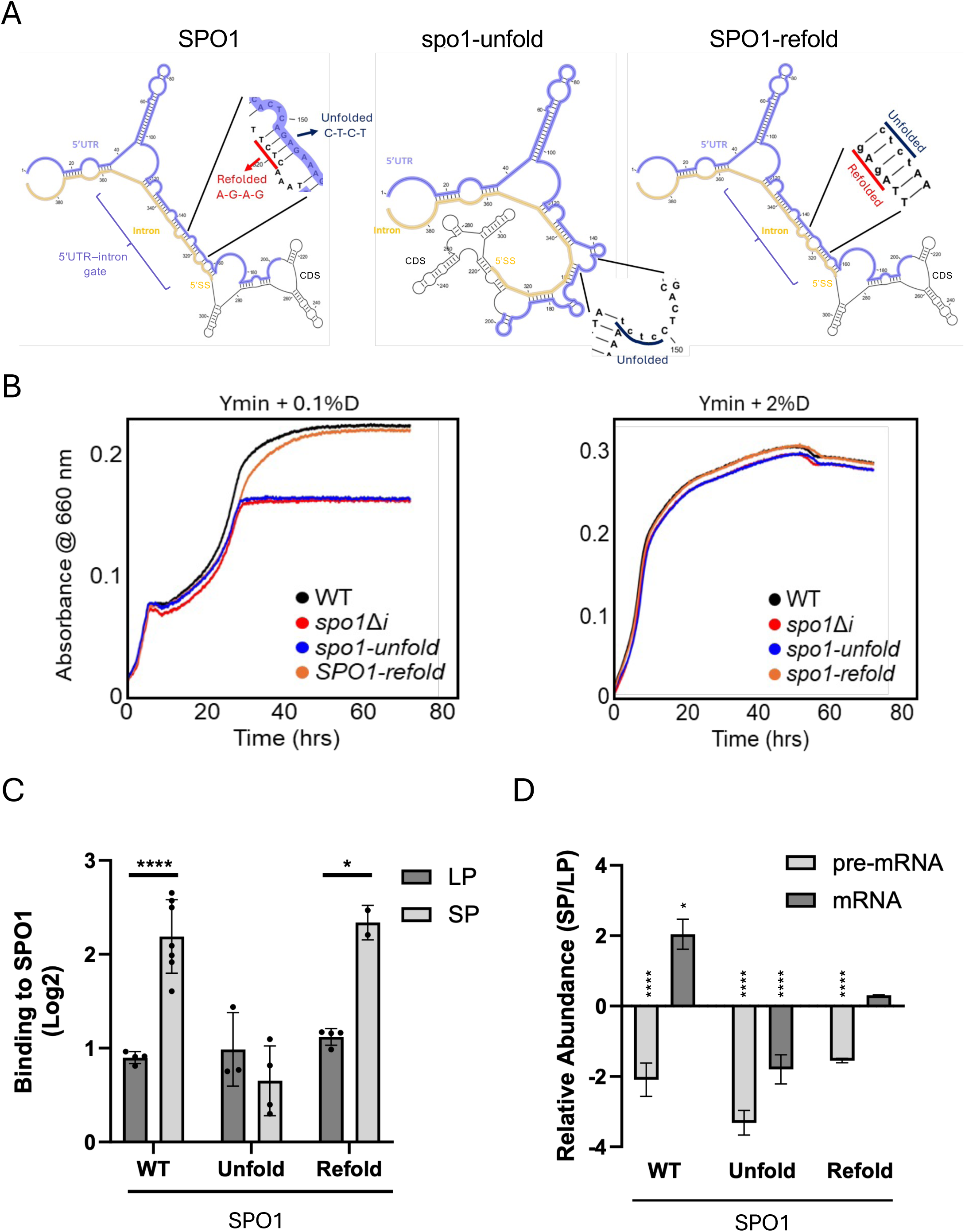
The 5′UTR–intron RNA gate drives starvation-dependent U1 snRNP association and splicing to support starvation fitness. **(A)** Structure-guided design of *SPO1* 5′UTR–intron gate mutants. The wild-type SPO1 transcript forms a predicted 5′UTR–intron structure near the 5′SS. The *spo1-unfold* mutant disrupts this pairing, whereas compensatory substitutions in *SPO1-refold* restore base pairing without restoring the wild-type primary sequence. **(B)** Growth of WT, *spo1Δi*, *spo1-unfold*, and *SPO1-refold* strains in amino acid-depleted minimal medium (Ymin) containing 0.1% or 2% dextrose. Growth was monitored by absorbance at 660 nm over 72h. Curves show the mean of at least three biological replicates. Disrupting the SPO1 gate phenocopies intron deletion under low-dextrose starvation conditions, whereas restoring base pairing rescues growth. **(C)** Starvation-induced U1 snRNP association with SPO1 RNA depends on the 5′UTR–intron gate. SPO1 RNA co-immunoprecipitated with Snu71-associated U1 snRNP in log phase (LP) and stationary phase (SP) was quantified by RT-qPCR, normalized to unaffected noncoding RNAs (RPR1 and NME1), and corrected by subtracting signal from an untagged control. Starvation increases U1 association with wild-type SPO1; this gain is lost in *spo1-unfold* and restored in *SPO1-refold*. Bars show mean ± SD from at least three biological replicates. Significance was determined using a two-sided t-test assuming unequal variances; *p<0.05; ****p<0.0001. **(D)** Starvation-induced SPO1 RNA processing depends on the 5′UTR–intron gate. Pre-mRNA and mature mRNA levels were quantified by RT-qPCR in cells grown in LP and after entry into SP, normalized to unaffected noncoding RNAs (RPR1 and NME1), and plotted as log2 SP/LP ratios (pre-mRNA, light gray; mature mRNA, dark gray). Starvation shifts wild-type *SPO1* output toward the mature mRNA state; this shift is impaired in *spo1-unfold* and restored in *SPO1-refold*. Bars show mean ± SD from three biological replicates. Significance between LP and SP was determined using a two-sided t-test assuming unequal variances; *p<0.05; ****p<0.0001.

### A synthetic gate is sufficient to reprogram a buffered DownSI transcript

We next tested whether a 5′SS-proximal 5′UTR–intron gate is sufficient to confer UpSI-like behavior on a DownSI transcript. *RPL18B* has a short 5′UTR and limited 5′SS-proximal 5′UTR– intron pairing, so we introduced a designed 5′UTR sequence that promotes pairing with the intron and mimics the UpSI gate (*RPL18B-fold*; Figure 4A). We hypothesized that the 5′UTR–intron gate is a transferable cis-regulatory element that can impose starvation-dependent U1 engagement and altered RNA processing on an otherwise buffered intron-containing transcript, and that installing such a gate in *RPL18B* would be sufficient to restore starvation growth in *spo1Δi* cells. At its native locus, the engineered gate did not impair growth: wild-type, *rpl18bΔi*, and chromosomal *RPL18B-fold* strains grew similarly in both 2% and 0.1% dextrose starvation media (Figure 4B, left). When expressed ectopically in *spo1Δi* cells, which also retain the native *RPL18B* locus, the plasmid-borne *pRPL18B-fold* rescued the low-dextrose growth defect (Figure 4B, right). Thus, a synthetic gate can stand in for *SPO1* intron function during starvation growth and reprogram a DownSI transcript into a starvation-responsive, U1-engaged state. This is consistent with the broader observation that it is the intron architecture, rather than the specific coding sequence, that is critical for starvation tolerance.

**Figure 4.**
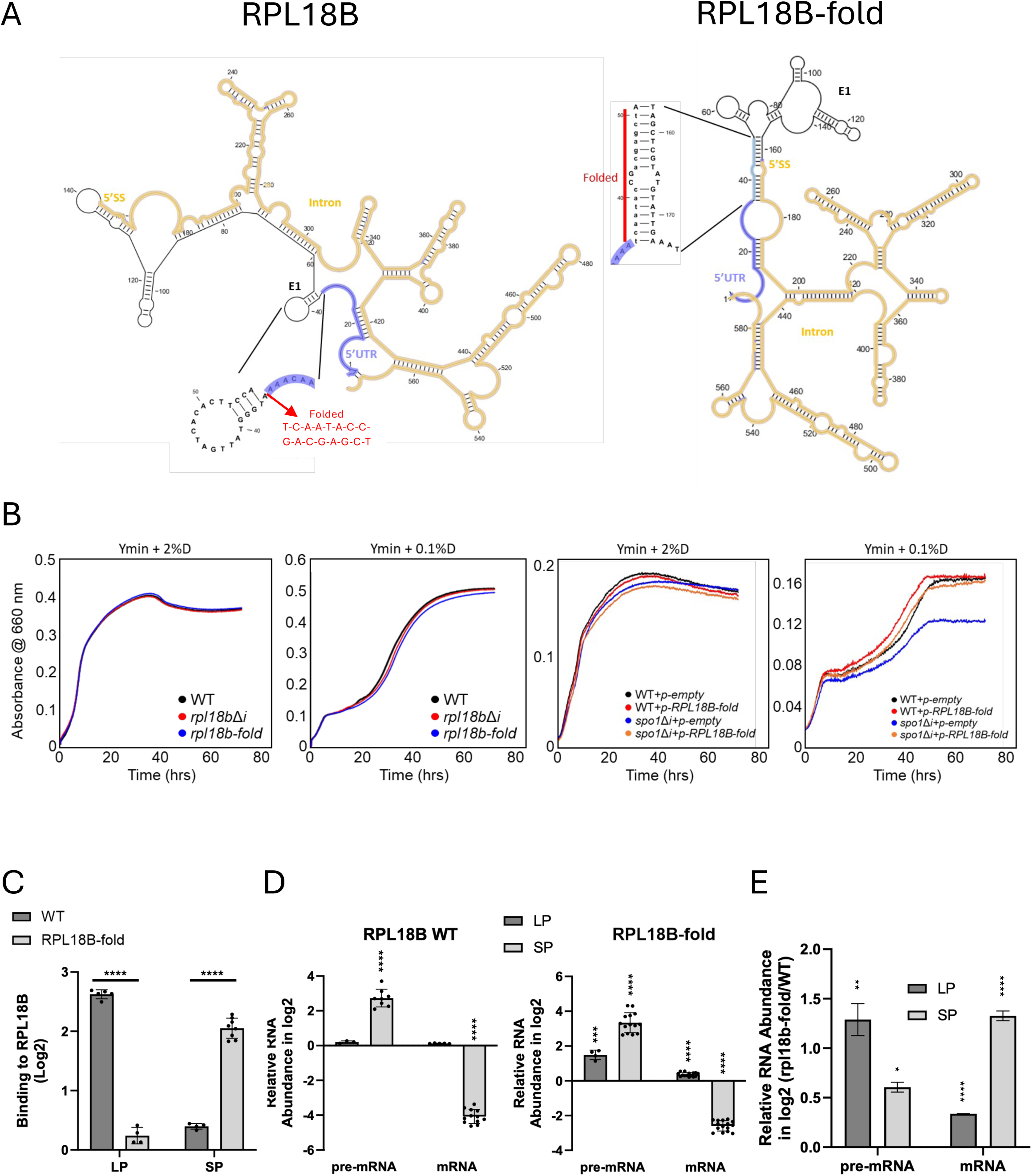
An engineered 5′UTR–intron RNA gate confers starvation-dependent U1 snRNP association and splicing and supports starvation growth. **(A)** Structure-guided design of an engineered RPL18B 5′UTR–intron gate. The wild-type RPL18B DownSI transcript forms limited 5′UTR–intron pairing, and the 5′SS region remains relatively accessible. In *RPL18B-fold*, sequence changes were introduced into the 5′UTR to promote base pairing with the intron and mimic the 5′SS-proximal 5′UTR–intron gate enriched in UpSIs. **(B)** Chromosomal and ectopic growth effects of the engineered RPL18B gate. Left, growth of WT, *rpl18bΔi*, and chromosomal *rpl18b-fold* strains in amino acid-depleted minimal medium containing 2% or 0.1% dextrose. These strains grow comparably, indicating that the engineered gate is not deleterious at the native *RPL18B* locus. Right, growth of WT or *spo1Δi* cells carrying empty vector or ectopic *pRPL18B-fold* in the same media. Growth was monitored by absorbance at 660 nm over 72 h, and curves show the mean of at least three biological replicates. Ectopic *RPL18B-fold* rescues the low-dextrose growth defect of *spo1Δi* cells. **(C)** The engineered gate confers starvation-induced U1 snRNP association on *RPL18B* RNA. *RPL18B* RNA co-immunoprecipitated with Snu71-associated U1 snRNP in log phase (LP) and stationary phase (SP) was quantified by RT-qPCR, normalized to unaffected noncoding RNAs (*RPR1* and *NME1*), and corrected by subtracting signal from an untagged control. Wild-type RPL18B shows little starvation-dependent change, whereas *RPL18B-fold* gains starvation-induced U1 association. Bars show mean ± SD from at least three biological replicates. Significance was determined using a two-sided t-test assuming unequal variances; ****p<0.0001. **(D)** The engineered gate confers starvation-dependent *RPL18B* RNA processing. Pre-mRNA and mature mRNA levels were quantified by RT-qPCR in LP (dark gray) and SP (light gray), normalized to unaffected noncoding RNAs (RPR1 and NME1), and plotted as log2 ratios for wild-type *RPL18B* and *RPL18B-fold*. Bars show mean ± SD from at least three biological replicates. Significance versus the wild-type *RPL18B* in LP was determined using a two-sided t-test assuming unequal variances; ***p<0.001; ****p<0.0001. **(E)** RNA abundance changes caused by the engineered *RPL18B* gate within each growth condition. Pre-mRNA and mature mRNA abundance are plotted as log2 *RPL18B-fold*/WT ratios in LP (dark gray) and SP (light gray). Bars show mean ± SD from at least three biological replicates. Significance between LP and SP was determined using a two-sided t-test assuming unequal variances; *p<0.05; **p<0.01; ****p<0.0001.

The engineered gate also changed U1 association. We validated TAP-Snu71 immunoprecipitation in the *RPL18B* backgrounds by western blot (Figure S4B). Wild-type *RPL18B* showed little starvation-induced U1 association, whereas *RPL18B*-*fold* gained U1 association in stationary phase (Figure 4C). This gain was not due to altered U1 snRNP recovery, because snR19 recovery was comparable between wild-type *RPL18B* and *RPL18B*-f*old* (Figure S4E). The synthetic structure therefore transfers a starvation-responsive U1 association pattern to a transcript that is normally buffered. This shift in U1 association was mirrored at the level of RNA output. Wild-type *RPL18B* accumulated pre-mRNA and lost mature mRNA in stationary phase, consistent with reduced splicing during nutrient depletion (Figure 4D). By contrast, *RPL18B*-fold dampened pre-mRNA accumulation and increased mature mRNA, most clearly under starvation (Figure 4D and 4E). Together, these data show that U1 recognition can be engineered through local pre-mRNA structure. By installing a 5′UTR–intron gate in *RPL18B*, we created starvation-induced U1 engagement, redirected RNA output toward mature mRNA, and restored growth when *SPO1* intron function was absent. Thus, the gate is not only required for *SPO1* regulation but is a portable RNA module capable of programming nutrient-controlled splice-site selection.

### Nam8 overexpression bypasses the RNA gate during starvation

Previous studies showed that yeast pre-mRNAs compete for limiting splicing machinery. Repression of abundant ribosomal protein gene (RPG) pre-mRNAs relieves this competition and increases splicing of non-RPG introns and protointrons^31,41^. Starvation also increases U1 abundance but does not increase U1 association with all transcripts. U1 association increases on introns whose splicing is induced by starvation and decreases on introns whose splicing is repressed^12^. Competition can therefore explain increased access to the spliceosome, but not selective U1 recruitment. We asked whether strengthening U1 contacts with intronic sequences downstream of the 5′ splice site could compensate for gate disruption. Nam8 contacts this region and stabilizes early U1–pre-mRNA complexes when 5′SS recognition is weak^42^. Nam8 also promotes meiotic splicing of introns with weak splice signals^43^. We therefore predicted that increasing Nam8 could bypass the requirement for gate pairing. As a control, we tested Luc7, a U1-associated factor that contacts the 5′ exon and stabilizes U1–pre-mRNA interactions^44^. *NAM8* overexpression rescued the low-dextrose growth defect of *spo1-unfold* but had no detectable effect in 2% dextrose (Figure 5A). *LUC7* overexpression did not rescue *spo1-unfold* in low dextrose and had no effect in 2% dextrose (Figure S5). Thus, Nam8-mediated rescue was specific to starvation and did not result from simply increasing any U1-associated protein.

**Figure 5.**
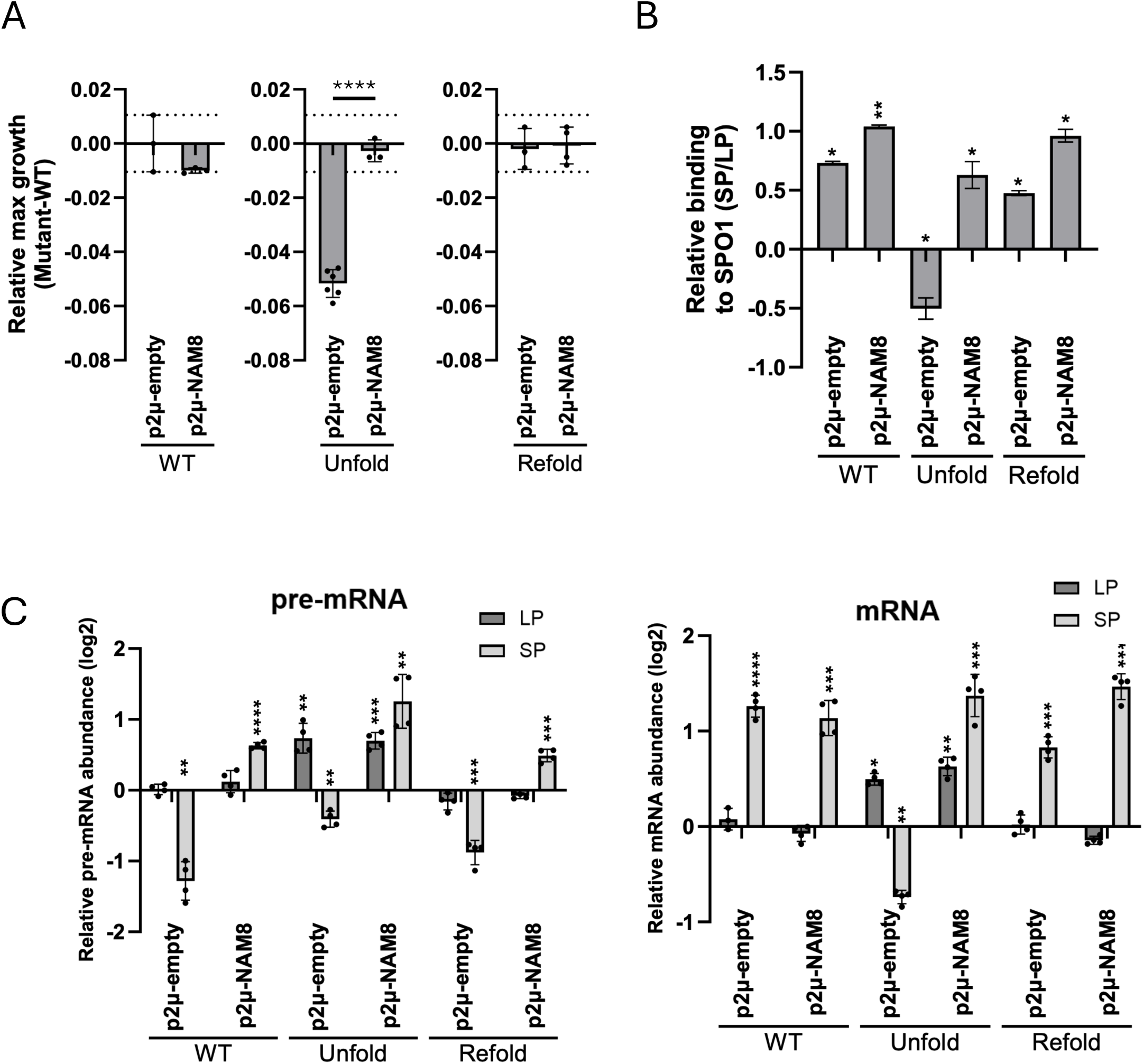
Nam8 sets the threshold of the 5′UTR–intron gate for U1 engagement and starvation fitness. **(A)** Nam8 bypasses the SPO1 gate defect to restore starvation fitness. Relative maximum growth is shown as each strain minus the WT strain containing the empty vector for WT, *spo1-unfold*, and *SPO1-refold* strains carrying empty vector or a *NAM8* overexpression plasmid in amino acid-depleted minimal medium containing 0.1% dextrose. Disruption of the *SPO1* 5′UTR–intron gate reduces starvation growth, whereas *NAM8* overexpression restores growth of *spo1-unfold* cells toward WT levels. WT and *SPO1-refold* cells show little additional benefit from *NAM8* overexpression, consistent with Nam8 acting preferentially when gate function is compromised. Bars show mean ± SD from at least three biological replicates; dotted lines indicate the SD range of WT controls. Significance was determined using a two-sided t-test assuming unequal variances where ****p<0.0001. **(B)** Nam8 reinstates U1 engagement with gate-disrupted SPO1 RNA. Relative U1 association with SPO1 RNA is shown as stationary phase/log phase (SP/LP) log2 ratios for WT, *spo1-unfold*, and *SPO1-refold* strains carrying empty vector or a *NAM8* overexpression plasmid. SPO1 RNA co-immunoprecipitated with Snu71-associated U1 snRNP was quantified by RT-qPCR, normalized to unaffected noncoding RNAs (RPR1 and NME1), and corrected by subtracting signal from an untagged control. Gate disruption reduces starvation-dependent U1 association, whereas *NAM8* overexpression restores U1 engagement with spo1-unfold RNA. Bars show mean ± SD from two biological replicates. Significance between LP and SP was determined using a two-sided t-test assuming unequal variances where *p<0.05 and **p<0.01. **(C)** Nam8 redirects gate-disrupted SPO1 RNA toward productive expression. Pre-mRNA and mature mRNA levels were quantified by RT-qPCR in log phase (LP) and stationary phase (SP) for WT, *spo1-unfold*, and *SPO1-refold* strains carrying empty vector or a *NAM8* overexpression plasmid. RNA abundance was normalized to unaffected noncoding RNAs (*RPR1* and *NME1*) and plotted relative to WT cells carrying empty vector in LP. *NAM8* overexpression shifts the RNA output of gate-disrupted *spo1-unfold* cells toward the starvation-associated expression pattern, linking restored U1 engagement to productive SPO1 RNA processing. Bars show mean ± SD from two biological replicates and two technical replicates. Statistical significance relative to WT cells carrying empty vector in LP was determined using a two-sided t-test assuming unequal variances; *p<0.05; ***p<0.001; ****p < 0.0001.

To determine whether this effect extended beyond *SPO1*, we examined *MMS2*. Disrupting the predicted *MMS2* gate impaired growth in low dextrose, whereas restoring the pairing improved growth. *NAM8*, but not *LUC7*, overexpression bypassed the gate-disrupted phenotype (Figures S6A–S6E). Nam8 can therefore compensate for gate disruption in two starvation-induced transcripts. Because growth rescue could occur indirectly, we tested whether Nam8 restored U1 association and RNA maturation. We confirmed TAP-Snu71 immunoprecipitation in the NAM8-overexpression strains by western blotting (Figure S4C). As shown above, *spo1-unfold* reduced starvation-dependent U1 association with *SPO1* RNA (Figures 3C and 5B). *NAM8* overexpression restored this association without changing the abundance of U1 snRNA, snR19 (Figures 5B and S4F–S4G). *NAM8* overexpression also increased mature *SPO1* RNA and shifted the RNA profile toward the mature state during starvation (Figure 5C). These results show that Nam8 compensates for disrupted gate pairing at the step of U1 recruitment. Thus, RNA structure and Nam8-dependent stabilization converge to promote selective 5′SS recognition during starvation.

### A starvation-induced RNA gate redirects U1 engagement to promote nutrient-stress adaptation

These findings explain how a change in pre-mRNA structure can alter splice-site choice during nutrient depletion. U1 binding at a 5’SS is not dictated by splice-site complementarity alone. It also depends on local transcript features and U1-associated proteins that help maintain the U1– pre-mRNA interaction^4^. In nutrient-replete cells, representative UpSI transcripts such as *SPO1* show limited proximal gate formation and low U1 snRNP association (Figure 6A). During starvation, the gate forms near the 5′ splice site, increases U1 interaction with the pre-mRNA and shifts RNA output toward mature mRNA (Figure 6B). DownSI transcripts, which have weaker or more distal 5′UTR–intron pairing relative to the 5′SS, remain buffered from this starvation-induced U1 response (Figure 6C). Three complementary experiments establish how the gate controls U1 engagement: disruption and restoration of SPO1 pairing, transfer of an engineered gate to *RPL18B*, and Nam8-mediated bypass of the disrupted SPO1 gate. Disrupting the *SPO1* gate reduces U1 association, impairs SPO1 RNA maturation, and compromises starvation growth, whereas restoring base pairing rescues each outcome (Figure 6D and 6E). Installing a synthetic gate in *RPL18B* creates starvation-induced U1 association and changes RNA output in a transcript that normally remains outside this response (Figure 6F). Increasing Nam8 provides a protein-based route to the same outcome: when the RNA gate is disrupted, Nam8 restores U1 association, *SPO1* RNA maturation, and starvation growth (Figure 6F). Together, these data show that starvation remodels local pre-mRNA structure to determine which transcripts are preferentially engaged by U1 during nutrient depletion. By directing U1 engagement toward introns that support growth under nutrient limitation, the RNA gate converts cellular state into splice-site choice, RNA maturation, and nutrient-stress adaptation.

**Figure 6.**
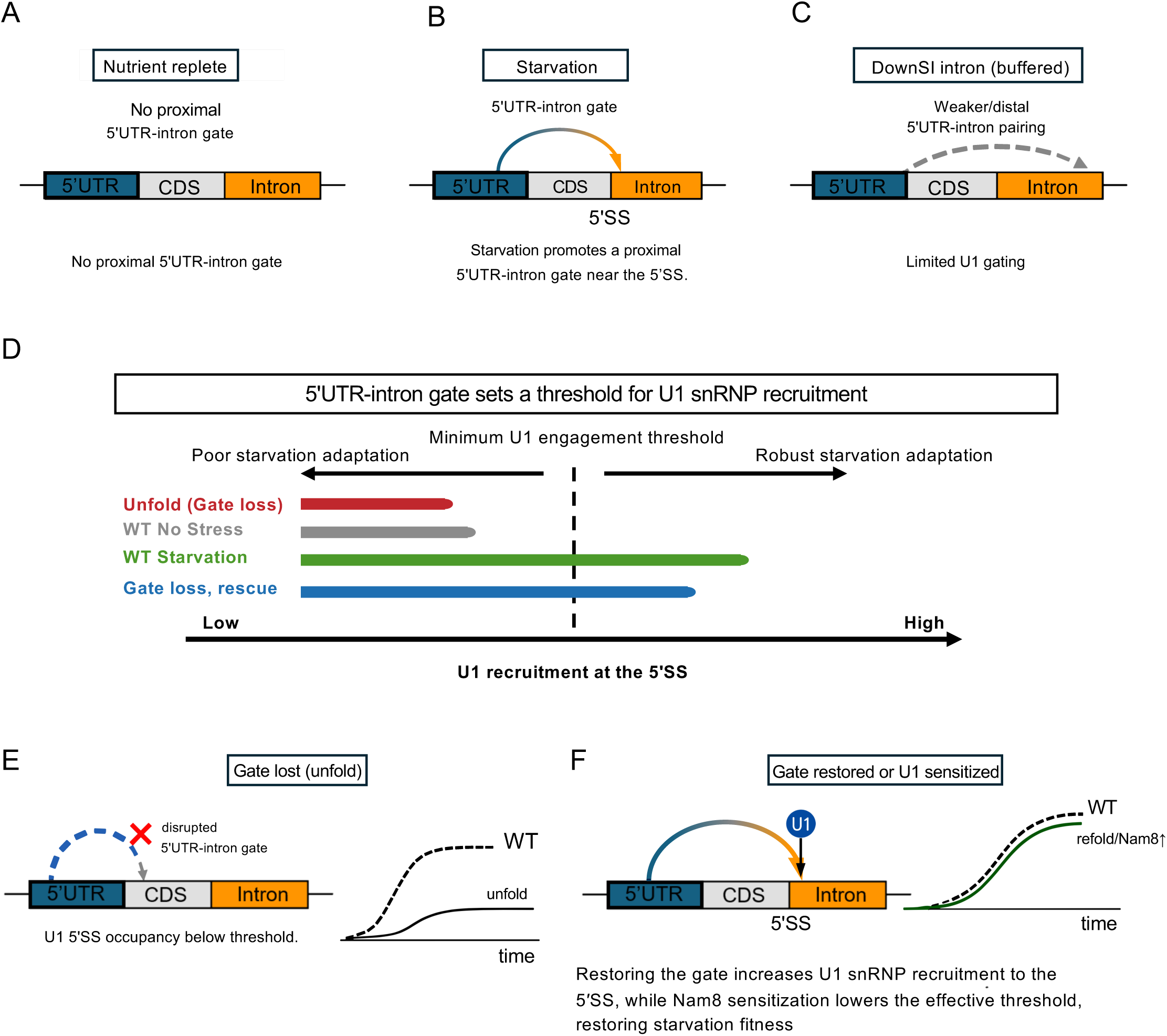
A starvation-induced 5′UTR–intron gate tunes U1 snRNP engagement to promote starvation fitness. **(A)** Model for a gate-dependent starvation-induced intron (UpSI) in nutrient-replete conditions. The 5′UTR–intron gate is weak or absent near the 5′ splice site (5′SS), leaving the 5′SS in a low-U1-engagement state. **(B)** During starvation, the same UpSI transcript remodels to form a proximal 5′UTR–intron gate near the 5′SS. This starvation-induced architecture promotes U1 snRNP engagement and supports productive RNA maturation. **(C)** Starvation-dispensable introns (DownSIs) are modeled as having weaker, more distal, or less responsive 5′UTR–intron pairing. As a result, these transcripts remain relatively buffered from starvation-induced U1 engagement and do not undergo the same gate-dependent increase in RNA processing. **(D)** The horizontal axis represents relative U1 engagement at the 5′SS, from low to high, based on the U1 RIP-qPCR, RNA-output, and growth data in Figures 3–5. The dashed line marks the minimum U1 engagement associated with productive RNA maturation and starvation adaptation. Gate loss (red), as in spo1-unfold, shifts the transcript below this threshold, reducing U1 association, RNA maturation, and starvation growth. In wild-type cells (green), starvation-induced gate formation increases U1 engagement to or above threshold in comparison to wild-type with no stress (gray), supporting productive maturation and growth. Nam8 overexpression (blue) compensates for gate loss by restoring U1 engagement above threshold, showing that the gate and Nam8 converge on the same step of selective U1 recognition rather than acting through independent mechanisms. **(E)** In the gate-disrupted spo1-unfold mutant, loss of 5′UTR–intron pairing reduces U1 engagement below the productive threshold. This below-threshold state is associated with impaired starvation-induced *SPO1* RNA maturation and reduced starvation fitness. **(F)** Restoring productive U1 engagement can occur through two routes. In the cis-acting route, compensatory refolding of the 5′UTR–intron gate increases U1 engagement at the 5′SS. In the trans-acting route, Nam8 overexpression lowers the effective requirement for the gate and restores U1-dependent RNA maturation despite gate disruption. Both routes return the transcript toward a productive U1-engaged state and support starvation growth toward the wild-type trajectory.

## DISCUSSION

U1 snRNP recognition of the 5′ splice site is an early commitment step in spliceosome assembly, but 5′SS sequence complementarity alone does not explain which introns are engaged in cells. Recent single-molecule and genetic studies support a model in which U1 binding is shaped by local RNA context, auxiliary U1 proteins, and cellular states^4,42^. Here, we identify a condition-responsive 5′UTR–intron RNA structure that provides this missing context during nutrient depletion. This 5′SS-proximal gate is enriched in starvation-induced introns, remodels *in vivo* during starvation, controls U1 association and RNA maturation, and can be engineered into a normally buffered transcript.

Our findings define the RNA-encoded mechanism underlying the nutrient-stress splice switch. Previous work showed that yeast introns promote growth under nutrient limitation and that starvation redistributes U1 snRNP occupancy, with many DownSI transcripts losing U1 association while UpSI transcripts gain U1 association and are preferentially engaged or processed^12,32^. What remained unknown was how individual pre-mRNAs encode this selectivity. We identify a starvation-remodeled 5′UTR–intron RNA gate positioned near, and not sequestering, the 5′SS as the cis-acting feature that distinguishes UpSIs from DownSIs. These findings establish local pre-mRNA architecture as a cis-acting determinant that converts nutrient state into selective, threshold-like U1 engagement, providing an additional regulatory layer beyond only simple spliceosome sponging or titration.

The gate model also explains why starvation does not simply cause global splicing repression. Previous work showed rapid, transcript-specific splicing changes during environmental stress and nutrient limitation, including repression of many ribosomal protein gene introns and persistence or accumulation of selected introns^12,32,45^. Our results add a mechanistic layer: starvation remodels the local architecture of specific pre-mRNAs so that selected introns remain competent for U1 engagement while others are buffered. In this model, DownSIs are not merely passive losers in a competition for spliceosomes; rather, they generally lack the proximal gate architecture that allows UpSIs to maintain or gain U1 association during stress.

The strongest evidence for the gate comes from separating RNA structure from primary sequence. Disrupting SPO1 gate pairing reduced U1 association, mature RNA output, and starvation growth, whereas compensatory mutations restored all three without restoring the wild-type sequence. Conversely, installing a 5′ splice-site-proximal gate in *RPL18B* converted this normally DownSI transcript into a starvation-responsive substrate, increasing U1 association and redirecting RNA output during starvation. These results indicate that the gate is not simply a marker of UpSIs but an instructive RNA element that can be moved to reprogram U1 engagement. The growth rescue by ectopic *RPL18B*-fold suggests that engineered gates can substitute for endogenous starvation-responsive introns.

Nam8 places the gate within the early U1-recognition pathway. Nam8 is a U1-associated RNA-binding protein that promotes splicing of weak or regulated introns and was originally identified as a multicopy suppressor of splicing defects caused by altered RNA folding^43,46,47^. Consistent with this role, *NAM8* overexpression restored U1 association, SPO1 RNA maturation, and starvation growth when the gate was disrupted, whereas *LUC7* overexpression did not. This specificity argues that Nam8 does not act as a general growth enhancer or nonspecific increase in U1-associated protein dosage. Instead, cis-acting gate structure and trans-acting Nam8 converge on the same limiting step: maintaining U1 interaction with selected pre-mRNAs during starvation by effectively lowering the U1 engagement requirements for gate-controlled splice site.

Previous studies have shown that direct base-pairing of the 5′SS within intronic structures can inhibit U1 recognition in vegetative growth^48^. Our gate model differs in two important aspects. First, the gate is formed by base pairing between the 5′UTR and the intron in a region adjacent to, rather than directly encompassing, the 5′SS, creating a distinct RNA architecture and folding dynamics rather than directly masking the U1-binding sequence. Second, unlike previously described inhibitory structures that appear constitutively repressive, the SPO1 gate functions in a condition-dependent manner: it reduces U1 engagement during vegetative growth but becomes permissive during starvation, when the spliceosome is remodeled and Nam8 promotes productive U1 recruitment. Thus, the novelty of our model is not simply the presence of an inhibitory RNA structure, but the discovery of a dynamic, 5′SS-proximal RNA gate whose effect on U1 recognition is rewired by the cellular state.

Several questions remain. We do not directly measure U1 dwell time, spliceosome commitment, or the physical geometry of the U1–gate complex, so the model is best stated as increased U1 association rather than proven kinetic stabilization. In addition, the predicted gate is enriched across UpSIs, whereas *in vivo* remodeling was tested in selected transcripts. Broader structure probing will be needed to define how widely this regulatory logic applies. UpSI gates also appear transcript-specific rather than structurally identical, suggesting that different folds may achieve a shared outcome: increased U1 engagement near the 5′SS under nutrient stress. Finally, how individual gate-controlled introns contribute to cellular adaptation remains to be resolved, especially because starvation splicing likely reflects both transcript-specific regulation and competition for limiting spliceosomal capacity.

The broader implication is that dynamic pre-mRNA structure can couple cellular physiology to splice-site choice. Although the specific *SPO1* gate and its functional relationship with Nam8 are unlikely to be conserved one-to-one, we propose that the underlying principle extends beyond yeast. In mammalian cells, local RNA structure influences the choice between competing 5′ splice sites ^49^, whereas TIA1 and TIAL1/TIAR perform a Nam8-like function by binding U-rich intronic sequences downstream of weak 5′ splice sites and promotingU1 recruitment through U1-C ^50,51^. These proteins also regulate disease-relevant splicing events, including inclusion of SMN2 exon 7 ^52^. Furthermore, small molecules can stabilize the U1–5′ splice-site interaction ^36^, whereas engineered U1 particles can be redirected to downstream intronic sequences to correct disease-associated splicing^53^. Together, these observations raise the possibility that condition-responsive RNA structures near mammalian splice sites could tune U1 recruitment and RNA output. Although a starvation-responsive gate analogous to the *SPO1* element has not been identified in mammals, our findings establish that such a gate can be required for adaptation, transferred to another transcript, and bypassed by increasing a U1-stabilizing factor. Dynamic pre-mRNA folding may therefore provide a general and programmable mechanism through which changes in cellular state are translated into selective splice-site recognition.

## RESOURCE AVAILABILITY

### Lead contact

Requests for further information and resources should be directed to and will be fulfilled by the lead contact, Sherif Abou Elela.

### Materials availability

All strains are available upon reasonable request except *Tap-SNU71* strain (obtained by material transfer agreement). Plasmids generated in this study will be shared with academic investigators subject to completion of a standard material transfer agreement, if required by the authors’ institution.

### Data availability

All data are available in the main text or the supplementary data. Additional data generated in this study have been submitted to the NCBI Gene Expression Omnibus (GEO; https://www.ncbi.nlm.nih.gov/geo) under the accession numbers GSE342187 (DMS-MaPseq) and will be publicly available as of the date of publication. Any additional information required to reanalyze the data reported in this paper is available from the lead contact upon reasonable request.

## ACKNOWLEDGEMENTS

We thank members of the Abou Elela, Rouskin, and Scott laboratories for discussions and critical feedback throughout this work. We also thank Dr Aaron Hoskins for the *Tap-SNU71* strain. This work was supported by the Canadian Institutes of Health Research (CIHR) [grant 413935 to S.A.] and the Research Chair in RNA Biology and Cancer Genomics [950-232264 to S.A]. Funding to pay for the Open Access publication charges for this article was provided by the Research Chair in RNA Biology and Cancer Genomics.

## AUTHOR CONTRIBUTIONS

Jasmine Tsang: Conceptualization, Formal analysis, Methodology, Visualisation, Writing— original draft. Julie Parenteau: Formal analysis, Visualisation, Writing—review & editing. Federico Fuchs Wightman and Kristina Song: Formal analysis, Methodology, Visualisation, Writing—review & editing. Silvi Rouskin and Michelle S. Scott: Writing—review & editing. Sherif Abou Elela: Conceptualization, Formal analysis, Methodology, Visualization, Funding acquisition, Writing—original draft.

## SUPPLEMENTAL INFORMATION

Supplementary Data are available at Molecular Cell online.

Supplementary Figures S1-S6.

Supplementary Tables S1-S2. Excel file containing primers used.

## STAR METHODS

KEY RESOURCES TABLE.

KEY RESOURCES TABLE
| REAGENT OR RESOURCE | SOURCE | IDENTIFIER |
| --- | --- | --- |
| Antibodies |  |  |
| TAP Tag Polyclonal Antibody | Invitrogen | Cat#CAB1001;<br>RRID:<br>AB_10709700 |
| PGK1 Monoclonal Antibody | Invitrogen | Cat#459250; RRID:<br>AB_2532235 |
| ECL Anti-Rabbit IgG | Cytiva | Cat#NA934V |
| ECL Anti-Mouse IgG | Cytiva | Cat#NA931V |
| Chemicals, peptides, and recombinant proteins |  |  |
| Dimethyl Sulfate | Sigma | Cat#D186309-100ml |
| Beta Mercaptoethanol | BioShop Canada | Cat#MER002.500 |
| Isoamyl Alcohol | VWR | Cat#97065-028 |
| Critical commercial assays |  |  |
| RNeasy Mini Kit | QIAGEN | Cat# 74104 |
| Deposited data |  |  |
| DMS-MapSeq Data | This paper | GEO: GSE342187 |
| Uncropped Western Blots (Figures S4A to S4C) | This paper | https://data.mendelley.com/preview/tpy68dtw83?a=a79d9513-435a-40fb-bc83-f83f276af923 |
| Experimental models: Organisms/strains |  |  |
| *S. cerevisiae*: Strain background: JPY10H3 (BY4741) | The Abou Elela Laboratory | N/A |
| Oligonucleotides |  |  |
| Primers for the construct of mutated strains, see Table S1 | This paper | N/A |
| Primers for PCR amplifications, see Table S2 | This paper | N/A |
| Software and algorithms |  |  |
| QIAGEN CLC Main WorkBench v.11.0.1 | QIAGEN | https://digitalinsights.qiagen.com/ |
| ViennaRNA package v.2.7.2 | Lorenz et al.<sup>54</sup> | https://link.springer.com/article/10.1186/1748-7188-6-26 |
| SEISMIC-RNA | Allan et al.<sup>55</sup> | https://www.biorxiv.org/content/10.1101/2024.04.29.591762v2 |

## METHODS DETAILS

### Strains and Plasmids

The *Tap*-Snu71 strain was synthesized as previously described^56^. The N-terminal fragment of the *SNU71* gene, including the *Tap* tag and the selection marker *HIS3MX6*, was amplified by PCR. *SPO1* and *RPL18B* with their intron removed (*spo1Δi* and *rpl18bΔi*) were obtained using the direct intron deletion strategy as previously described^12^. *SPO1* and *RPL18B* mutant constructs were generated by introducing defined sequence substitutions within regulatory regions relative to the start codon. For *SPO1*, mutations were introduced at two sites: the sequence CACTCAGAGAAA (−49 to −53 relative to the start codon) was replaced with CACTCCTCTAAA for the unfold, and AGTTAAACTCTT (+109 to +113 relative to the start codon) was replaced with AGTTAGAGTCTT for the refold. For *RPL18B*, an additional sequence AACAAATCAATACCGACGAGCTATGG was added just before the start codon (–1). The PCR products were digested with NotI and BamHI (*SPO1*) or XhoI and BamHI (*RPL18B*) and cloned into pRS316. *MMS2* strains were synthesized as previously described^32^. To amplify yeast *NAM8* and *LUC7* genes from wild-type genomic DNA, we used primers (Supplementary Table 1) with restriction sites; HindIII and SpeI for *NAM8* (–428 to +2059 nucleotides relative to the initiation codon) or XhoI and BamHI for *LUC7* (–510 to +1044 nucleotides relative to the initiation codon). The PCR products were digested with respective enzymes and cloned into 2-micron plasmid pRS426 (*NAM8*) or pRS423 (*LUC7*) to generate p2µ-NAM8 and p2µ-LUC7. The different strains: WT, *spo1Δi*, *spo1-unfold*, *SPO1-refold*, mms2*Δi, mms2-unfold*, *MMS2-refold* were then transformed with the 2-micron NAM8 or LUC7 plasmid. Yeast cells were transformed using a lithium acetate / single-stranded carrier DNA / polyethylene glycol method as described previously^57^. Transformants were grown in selective minimal medium lacking uracil for pRS426 plasmids or histidine for TAP-tagged strains and pRS423 plasmids. All plasmids were propagated in *Escherichia coli* strain using standard strains and conditions^58^. All primers used in this study to construct the strains are listed in Supplementary Table S1.

### Growth assays

Synthetic minimal media (1.7 g/L of yeast nitrogen base without amino acids, ammonium sulfate, and 1g/L of L-glutamic acid supplemented with amino acids to meet auxotrophic requirements) enriched with 0.1% and 2% D-glucose were used for yeast growth assays. Plasmids were grown in minimal media with the different levels of glucose without uracil (pRS426) or histidine (pRS423). Cells grown to the logarithmic phase were harvested by centrifugation and resuspended at a concentration of 3.36 × 10⁷ cells/mL in 1 mL of Ymin + 2%D. 5 µl of the cell suspension were diluted in 95 µl of Ymin (without or with 2% dextrose) in 96-well polystyrene microplates. The growth was monitored in microplate readers (Epoch2 or SynergyHTX) at 30°C with linear agitation for 72 hours, with optical density readings taken at 10-minute intervals. Maximum growth is determined after 72 hours of growth. All growth assays were performed with at least three biological samples and two to three technical replicates.

### Secondary structure prediction

Secondary structures were modeled using QIAGEN CLC Main WorkBench v.11.0.1 (QIAGEN Digital Insights Bioinforma. Softw. QIAGEN Digit. Insights. https://digitalinsights.qiagen.com/) secondary structure analysis tool (build 1305231204) based on gene sequences obtained from the Saccharomyces Genome Database (SGD)^59^. The structures were predicted using the WorkBench platform, which employs a modified version of the algorithm developed by Michael Zuker^60^. The prediction of secondary structures is based on a free energy minimization approach that identifies the most thermodynamically stable conformation. The stability of the structure’s formation is defined by the amount of energy released. The more energy released, the higher the probability of structure forming.

### RNA secondary structure base-pairing analysis

RNA secondary structure predictions were performed using RNAfold –p from the ViennaRNA package (v2.7.2)^54^ to compute the base pair probabilities. Input sequences comprised the first exon (5’UTR and beginning of the CDS) and intron of each gene from the *S. cerevisiae* genome (R64-1-1, Ensembl release 115). Only base pairs forming a consecutive stem of ≥3 stacked pairs with probability of ≥0.5 were retained. For visualization, each gene’s sequence coordinates were normalized to a common display space anchored at the 5’ splice site with region widths scaled to the group median length of each region (separately for UpSI and DownSI genes). Base-pairing contacts were clustered across genes using KMeans with the gap statistic to determine the optimal number of clusters k (maximum k=10). Clusters supported by fewer than 15% of genes within a group were excluded. The median branch point position (show in red in Figure 1C) was computed as the median of per-gene branch point fractions (branch point position/intron length). To assess cross-species conservation of 5’UTR-intron base-pairing contacts, orthologous 5’UTR, beginning of CDS (together constituting the first exon) and intron sequences for UpSI and DownSI genes were retrieved from NCBI genome assemblies for nine yeast species: *S. paradoxus* (GCA_002079055.1), *S. kudriavzevii* (GCA_947243775.1), *S. mikatae* (GCA_947241705.1), *S. cerevisiae* (GCA_000146045.2), *S. castellii* (GCA_000237345.1), *S. barnettii* (GCA_903064755.1), *S. dairenensis* (GCA_000227115.2), *S. unisporus* (GCA_964275045.1) and *S. servazzii* (GCA_964275055.1). RNA secondary structure predictions were performed independently for each species sequence as described above.

The linear regression analyses, Pearson’s correlation coefficients, and corresponding P-values were calculated using GraphPad Prism (version 11.0.2) or Statistics Kingdom web page (https://www.statskingdom.com/linear-regression-calculator.html).

### *In vivo* DMS treatment and RNA extraction

Yeast cultures were grown in minimal media with 2% D-glucose until an OD_600_ of 0.6 was reached (logarithmic phase, LP) or for an additional 48 hours following the attainment of the logarithmic phase (stationary phase, SP). For the SP cultures, they were adjusted to contain the same number of cells than the LP cultures and diluted to the same final volume. For each condition, three biological replicates were subjected to *in vivo* dimethyl sulfate (DMS) treatment, and three biological replicates did not receive treatment. Cells were incubated with DMS at a final concentration of 7.5% (v/v) for 7 min at 30°C with vigorous mixing in a thermomixer. Reactions were quenched by adding two volumes of ice-cold stop solution containing β-mercaptoethanol (BME), isoamyl alcohol, and DEPC-treated water (9:15:6, v/v/v). Samples were mixed thoroughly, placed on ice, and centrifuged at approximately 3500g for 4 min at 4°C. Cell pellets were washed with wash solution (3:7 BME:DEPC-treated water), centrifuged again under the same conditions, and the supernatant was discarded. Total RNA was extracted using a hot acid phenol protocol. Cell pellets were resuspended in RNA lysis buffer (0.5M EDTA pH 8.0, 3M sodium acetate, and DEPC-treated water). SDS was added to a final concentration of 1.25% (w/v), and samples were incubated at 65°C for 3 min. Lysates were transferred to preheated acid phenol (pH 4.5) and incubated in the same conditions with vigorous mixing. Samples were rapidly frozen in a dry ice/ethanol bath and centrifuged. The aqueous phase was recovered and extracted a second time. RNA was precipitated, pelleted and washed with 80% ethanol, air-dried, and resuspended in 10 mM Tris-HCl (pH 7.0). RNA samples were then treated with TURBO DNase (Invitrogen, AM2238). Samples were incubated at 37°C for 20 min and subsequently purified using the Monarch Spin RNA Cleanup Kit (500 µg; New England Biolabs, T2050L) according to the manufacturer’s instructions, with elution in nuclease-free water.

### DMS-MaPseq reverse transcription and PCR

Reverse transcription was performed using Induro Reverse Transcriptase (New England Biolabs, M0681L). For each reaction, RNA was combined with primers (10µM). Samples were denatured at 65°C and immediately chilled on ice. Reverse transcription was carried out at 25°C for 5 min followed by 55°C for 45 min. Subsequently, 4M NaOH was added and reactions were heated at 95°C. cDNA products were purified using the Monarch Spin PCR & DNA Cleanup Kit (New England Biolabs, T1130L) and eluted in nuclease-free water. Target cDNAs were amplified using Q5 Hot Start High-Fidelity 2X Master Mix (New England Biolabs, M0494L). PCR conditions consisted of an initial denaturation at 98°C for 30s; 28 cycles of 98°C for 10s, 60°C for 10s, and 72°C for 10s; followed by a final extension at 72°C for 5 min. Amplicons were verified by electrophoresis on agarose gels stained with Safe DNA Gel Stain (Apex Bio, A8743). PCR products were purified using a double-sided SPRI bead size-selection procedure. Samples were first incubated with 0.55X bead volume, and the supernatant was subsequently subjected to a second purification with an additional bead volume to capture the desired DNA fragments. Beads were washed twice with 80% ethanol, and DNA was eluted in nuclease-free water.

### DMS-MaPseq Illumina library preparation and sequencing

Sequencing libraries were prepared from 200ng purified PCR product using the NEBNext UltraExpress DNA Library Prep Kit (New England Biolabs, E3325L) according to the manufacturer’s instructions. Library enrichment was performed using five PCR cycles. Sample barcoding was achieved using NEBNext Multiplex Oligos for Illumina (New England Biolabs, E6440L). Indexed libraries were sequenced on either an Illumina NextSeq 1000 platform or an iSeq 100 using a NextSeq 1000/2000 P1 XLEAP-SBS Reagent Kit (300 cycles; Illumina, 20100982) or an iSeq 100 i1 Reagent v2 (300 cycles; Illumina, 20031371), respectively. Paired– end sequencing was performed with 151 cycles for Read 1 and Read 2 and 8 cycles for each index read. Demultiplexing was carried out using Illumina BaseSpace.

### DMS-MaPseq analysis

Demultiplexed FASTQ files were processed using SEISMIC-RNA^55^. Quality control, adapter trimming, alignment, mutation calling, and RNA structure inference were performed using the default workflow (wf command) with the “--fold” and “--export” options enabled. Additional parameters were applied for the SPO1 amplicon (“--no-fastp-detect-adapter-for-pe”, “--fastp-adapter-1 AGATCGGAAGAGCACACGTCTGAACTCCAGTCA”, “--fastp-adapter-2 AGATCGGAAGAGCGTCGTGTAGGGAAAGAGTGT”, “--keep-discontig”, “--min-mut-gap=0”, “--min-mapq 40”, and “--min-finfo-read 0.99”) to improve adapter handling and alignment accuracy. Resulting datasets were inspected using SEISMICgraph ^61^ to assess alignment quality, sequencing coverage, mutation frequencies, and reproducibility between biological replicates. DMS-MaPseq quality control metrics are presented in Supplementary Figure S3A-D.

### Data visualization and structural analyses

Data visualization and statistical analyses were performed in R using the ggplot2 and patchwork packages. DMS reactivities were calculated as population-average mutation frequencies at adenine and cytosine residues. Primer-binding regions were excluded from all analyses, and an additional low-coverage region (positions 145–165) was masked in the SPO1 amplicon. Arc plots were generated from SEISMIC-RNA folding ensembles by selecting, for each condition, the structure exhibiting the highest median base-pair Jaccard similarity across replicates. Base pairs were classified as shared, LP specific, or SP specific. Local similarity between conditions was assessed by calculating Pearson correlation coefficients for DMS reactivities within 45-nt sliding windows containing at least five informative nucleotides.

### Cell culture and crosslinking

The labeled and unlabeled yeast strains were cultured in a minimal medium (+*lys, his, ura,* and *leu*) with 2% D-glucose until an OD_660_ of 0.4–0.5 was reached (LP) or for an additional 48 hours following the attainment of the logarithmic phase (SP). Formaldehyde (1% v/v) was added to the cell culture for crosslinking for 20 minutes at room temperature, with the medium stirred periodically. This reaction was inhibited with 0.36 M glycine for 5 minutes. The cells were then washed twice with cold TBS (20 mM Tris-HCl, pH 7.5, 150 mM NaCl) and aliquoted before being stored at –80°C.

### Immunoprecipitation

The cell pellets are resuspended in FA lysis buffer (50 mM HEPES-KOH, pH 7.5, 150 mM NaCl, 1 mM EDTA, 1% Triton X-100, 0.1% sodium deoxycholate, protease inhibitors) for LP cell pellets or in a lysis buffer (50 mM Tris-HCl, pH 7.5, 150 mM NaCl, 15% glycerol, 0.5% Tween 20, protease inhibitor) for SP pellets. Cells were lysed with glass beads in a Precellys homogenizer (Laboratory Supply Network) for six 10-second cycles at 6,500 rpm (LP) or four 10-second cycles at 6,500 rpm (SP). Following lysis, Branson sonication (8 pulses of 10 seconds at 20% amplitude) was performed to fragment the chromatin before centrifuging the cell extracts at 13,000 rpm for 15 minutes at 4°C. The supernatant was treated with DNase (0.1 mg/mL; Sigma) along with 5 mM CaCl₂, 25 mM MgCl₂, and 100 U/mL RNase Out (Invitrogen) for 20 minutes at 37°C. The lysates, with or without labels, are incubated for 16 hours at 4°C with magnetic beads (Dynabeads^TM^ Pan Mouse IgG, Life Technologies) without antibodies in TBS containing 5 mg/ml bovine serum albumin (BSA). The beads are washed sequentially with FA buffer (twice, for 3 minutes), FA buffer with 500 mM NaCl, and a wash buffer (10 mM Tris-HCl, pH 8, 250 mM LiCl, 1 mM ethylenediaminetetraacetic acid, 0.5% NP-40, 0.5% sodium deoxycholate) at 4°C. The specific enrichment of TAP-Snu71 was confirmed by Western blot. The immunoprecipitated complexes are eluted and treated with proteinase K (400 µg/mL in 1% SDS-TE buffer) for 2 hours at 42°C, followed by denaturation of the cross-links with a 1-hour incubation at 65°C. The immunoprecipitated RNA is reverse-transcribed for analysis by RT-qPCR. To assess RNA enrichment, the data are normalized against control samples (input) and against RPR1 and NME1, two non-coding RNAs, as an internal control. The enrichment calculation is as follows:

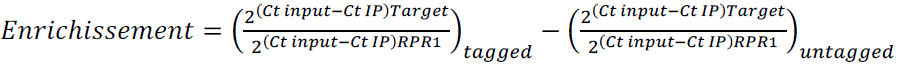

where Ct represents the threshold cycle value for each sample. The RNA enrichment represents the average of three biological samples, and the oligonucleotide sequences are listed in Supplementary Table S2.

### RNA extraction and RT-qPCR

Total yeast RNA is isolated from cell cultures grown in minimal medium containing 2% D-glucose during the logarithmic/exponential phase, when the OD₆₆₀ reaches approximately 0.4, or until the stationary phase is reached 48 hours after the logarithmic phase, using hot acid-phenol extraction. Only RNA samples with an A260/A280 ratio between 1.8 and 2.0 were used for RT-qPCR. The samples were treated with DNase using the Qiagen RNase-Free DNase Kit before using 50 ng for reverse transcription (RT) with MMuLV Reverse Transcriptase (MRT) from the University of Sherbrooke’s protein purification platform, a random p(dN)6 primer, dNTPs, and RNase Out. Each 10 µl of qPCR sample contains 3 µl of complementary DNA, 5 µl of SYBR Green Master Mix, and 2 µl of primers. Using a Bio-Rad C1000 Touch thermal cycler, the amplification cycle is as follows: 1 minute at 95°C; 50 cycles of 10 seconds at 95°C, 15 seconds at 60°C, 15 seconds at 72°C; 5 seconds at 65°C, and 5 seconds at 95°C. Each assay includes at least three biological replicates and two technical replicates. Transcripts are normalized using non-spliceosome RNAs (SPT15, ARO3, NME1, and RPR1). Relative expression is calculated using the 2^-ΔΔCt method. The oligonucleotides used for qPCR are described in Supplementary Table S2.

### Western blot analysis

Protein extracts (30 µg/well) quantified using the Bradford assay, and the BLUelf prestained protein ladder (Froggabio Inc.) were separated by 10% sodium dodecyl sulfate polyacrylamide gel electrophoresis (SDS-PAGE), followed by transfer to a Protran (GE Healthcare). The nitrocellulose membranes were blocked with a TBS-T solution (20 mM Tris-HCl, pH 7.6, 150 mM NaCl, 0.1% Tween 20) containing 5% skim milk overnight at 4°C. The membranes were incubated with the primary antibodies (rabbit anti-TAP #CAB1001 from ThermoFisher Scientific, 1:1000; mouse anti-Pgk1 #459250 from Invitrogen, 1:10,000) diluted in TBS-T with 5% skim milk for 90 minutes at room temperature. Following four 10-minute washes with TBS-T, a second incubation is performed with HRP-conjugated secondary antibodies (donkey anti-rabbit IgG #NA934V from GE Healthcare, 1:2000; sheep anti-mouse IgG #NA931V from GE Healthcare, 1:2000) diluted in TBS-T with 5% skim milk for 90 minutes at room temperature. After another four washes with TBS-T, detection was performed using Clarity Western ECL (Bio-Rad). The images obtained using a LAS4000 system (GE Healthcare) were then quantified with Bio-Rad’s Quantity One Software. At least three biological replicates of each condition were included.

