## Supplemental Figures for "A starvation-remodeled pre-mRNA structure controls U1 recruitment and nutrient-stress adaptation in yeast"

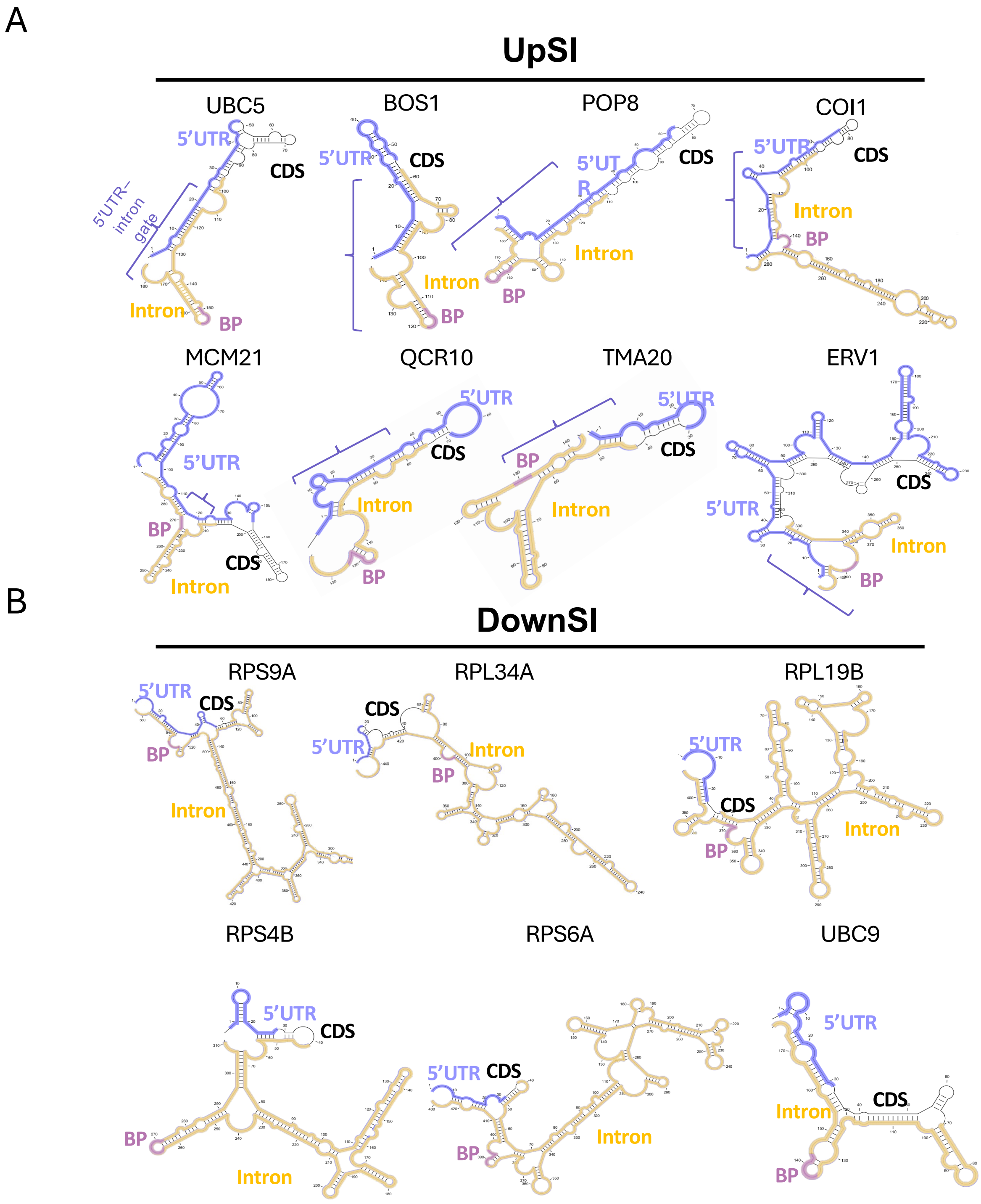


**Figure S1. Predicted minimum-free-energy secondary structures of representative UpSI and DownSI transcripts, related to Figure 1.** Predicted minimum-free-energy secondary structures spanning the 5′UTR (blue), coding sequence (gray), intron (yellow), and branch point (BP, purple) for representative starvation-induced introns (UpSIs) (**A**) and starvation-dispensable introns (DownSIs) (**B**). Structures were computed from the 5′UTR–CDS–intron region of each transcript using CLC Main Workbench with default parameters. (**A**) UpSI transcripts UBC5, BOS1, POP8, COI1, MCM21, QCR10, TMA20, and ERV1. (**B**) DownSI transcripts RPS9A, RPL34A, RPL19B, RPS4B, RPS6A, and UBC9. The 5′ splice site (5′SS) and the 5′UTR–intron gate are indicated where resolved. Consistent with the conserved signature in Figure 1, UpSI transcripts form extensive 5′UTR–intron pairing that folds back over the 5′SS-proximal region, whereas DownSI transcripts show more distal and heterogeneous 5′UTR–intron contacts with a comparatively accessible 5′SS.

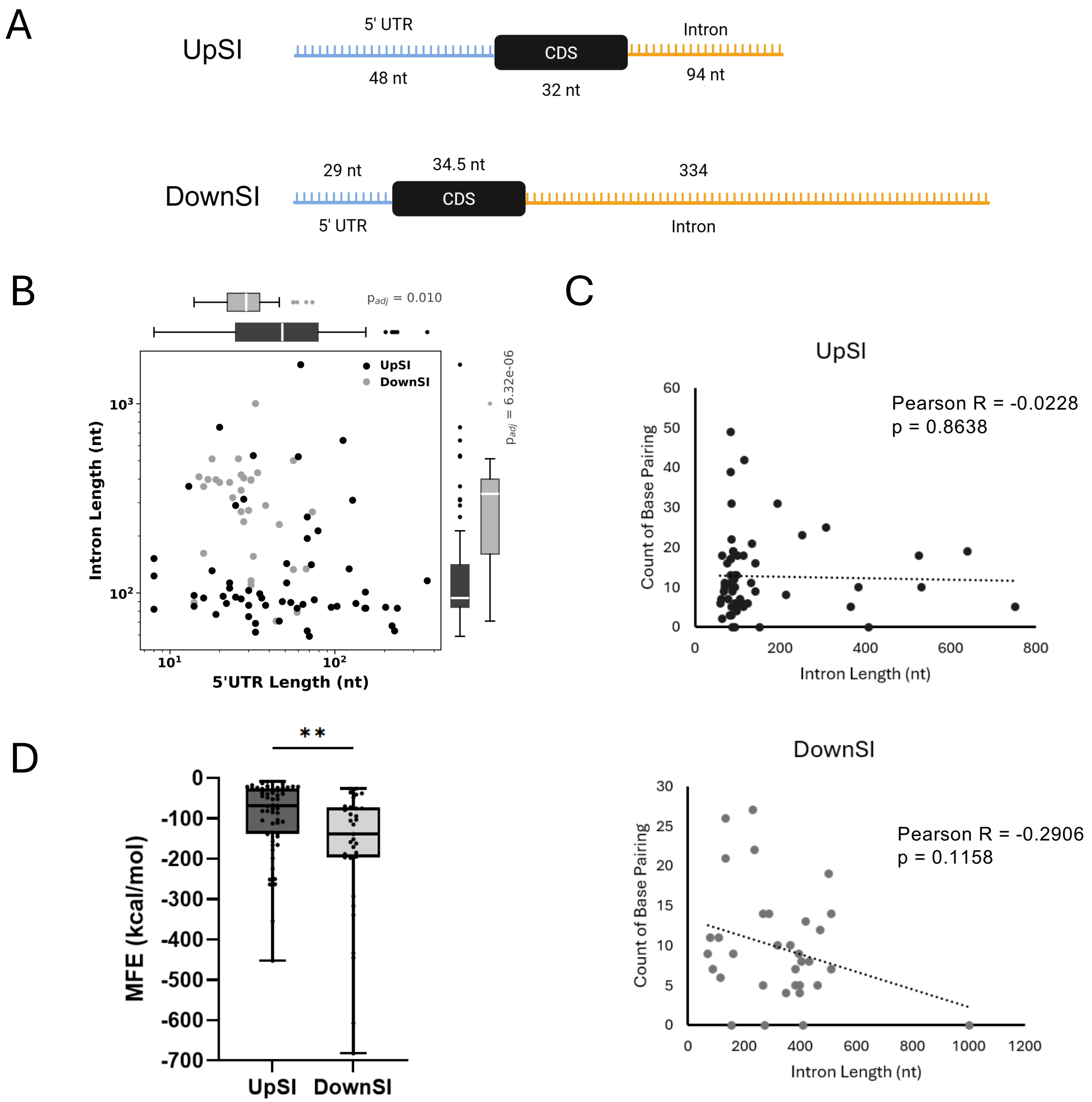


**Figure S2. Starvation-induced and starvation-dispensable introns differ in 5′UTR length, intron length, and predicted structural stability, related to Figure 1.** (**A**) Transcript architecture of UpSI and DownSI genes. Left, schematic comparing median 5′UTR (blue), CDS (gray), and intron (yellow) lengths between UpSI (5′UTR 48 nt, CDS 32 nt, intron 94 nt) and DownSI (5′UTR 29 nt, CDS 34.5 nt, intron 334 nt) genes. (**B**) Relationship between 5′UTR length and intron length across intron classes. Scatter plot of 5′UTR length versus intron length for UpSI (black) and DownSI (gray) genes; each dot represents one gene. Marginal box plots show the distribution of 5′UTR length (top) and intron length (right) for each class. Significance was determined using the Mann-Whitney U test with Benjamini-Hochberg correction; reported p values are $p = 0.010$ (5′UTR) and $p = 8.32 \times 10^{-6}$ (intron). **(C)** Relationship between the intron length and count of base pairs in the UpSIs (top) and DownSIs (bottom) where each dot represent a gene. Significance and corresponding correlation coefficients were determined by the Pearson test. (**D**) Predicted thermodynamic stability of UpSI and DownSI structures. Box plots of the minimum free energy (MFE) of predicted secondary structures for UpSI and DownSI transcripts, computed with RNAfold. UpSI structures have a higher (less negative) MFE than DownSI structures, indicating lower predicted thermodynamic stability. Boxes show the median and interquartile range, and whiskers show the full range. Significance was determined using Welch's t test (two asterisks, $p < 0.01$).

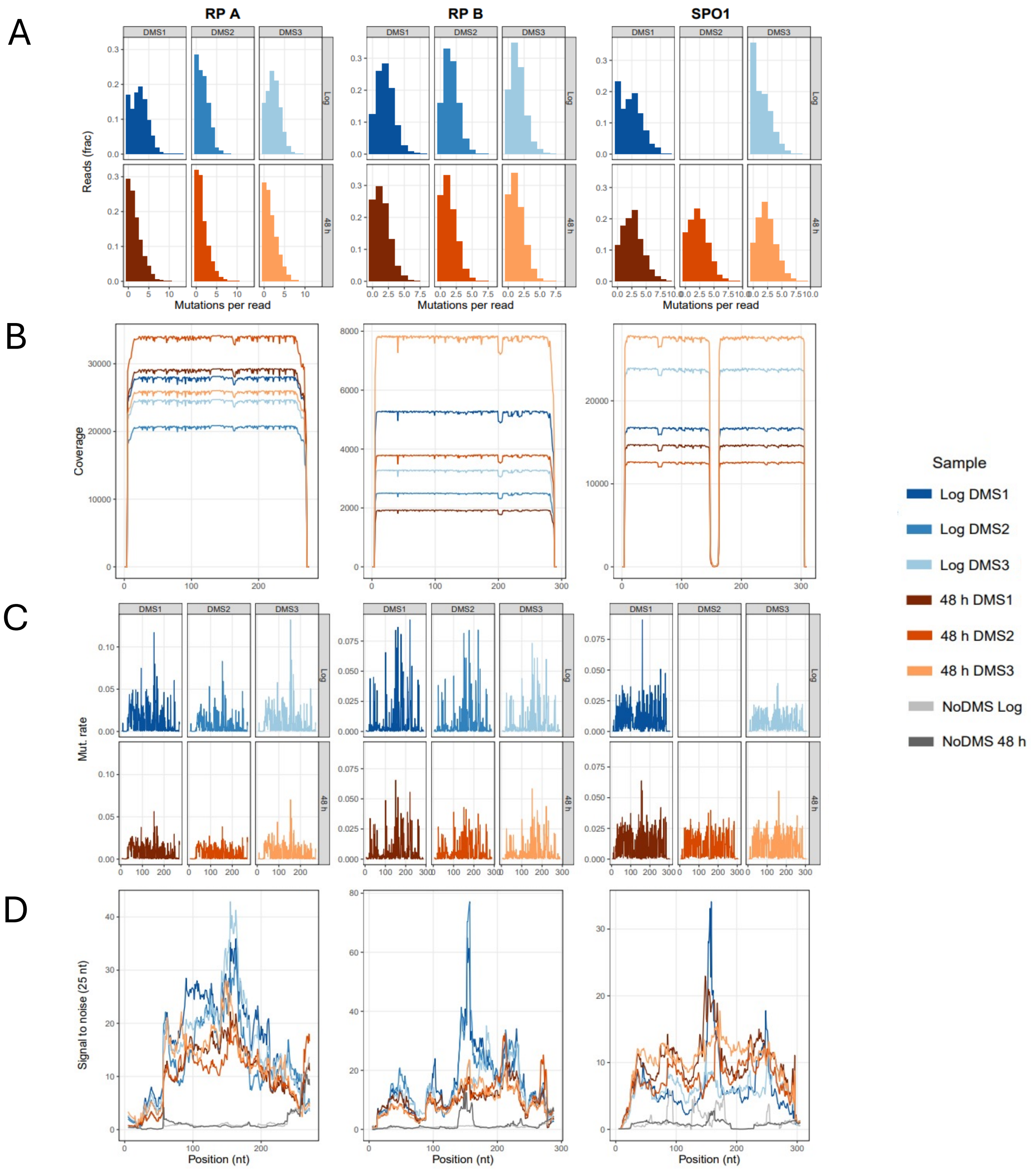


**Figure S3. DMS-MaPseq quality control, related to Figure 2.** (**A**) Distribution of per-read mutation counts for DMS-treated samples used for in vivo DMS-MaPseq analysis. (**B**) Read coverage across each amplified transcript region. RPL18B was analyzed as two amplicons, RPL18B_A and RPL18B_B, which were concatenated for downstream structural analysis. (**C**) Raw DMS reactivity profiles, calculated as the fraction of reads carrying a mutation at each nucleotide position. (**D**) Rolling signal-to-noise ratio across each transcript region using a 25-nt window. Signal-to-noise was calculated as the average reactivity of adenines and cytosines divided by the average reactivity of guanines and uracils. No-DMS controls are shown as background reference.

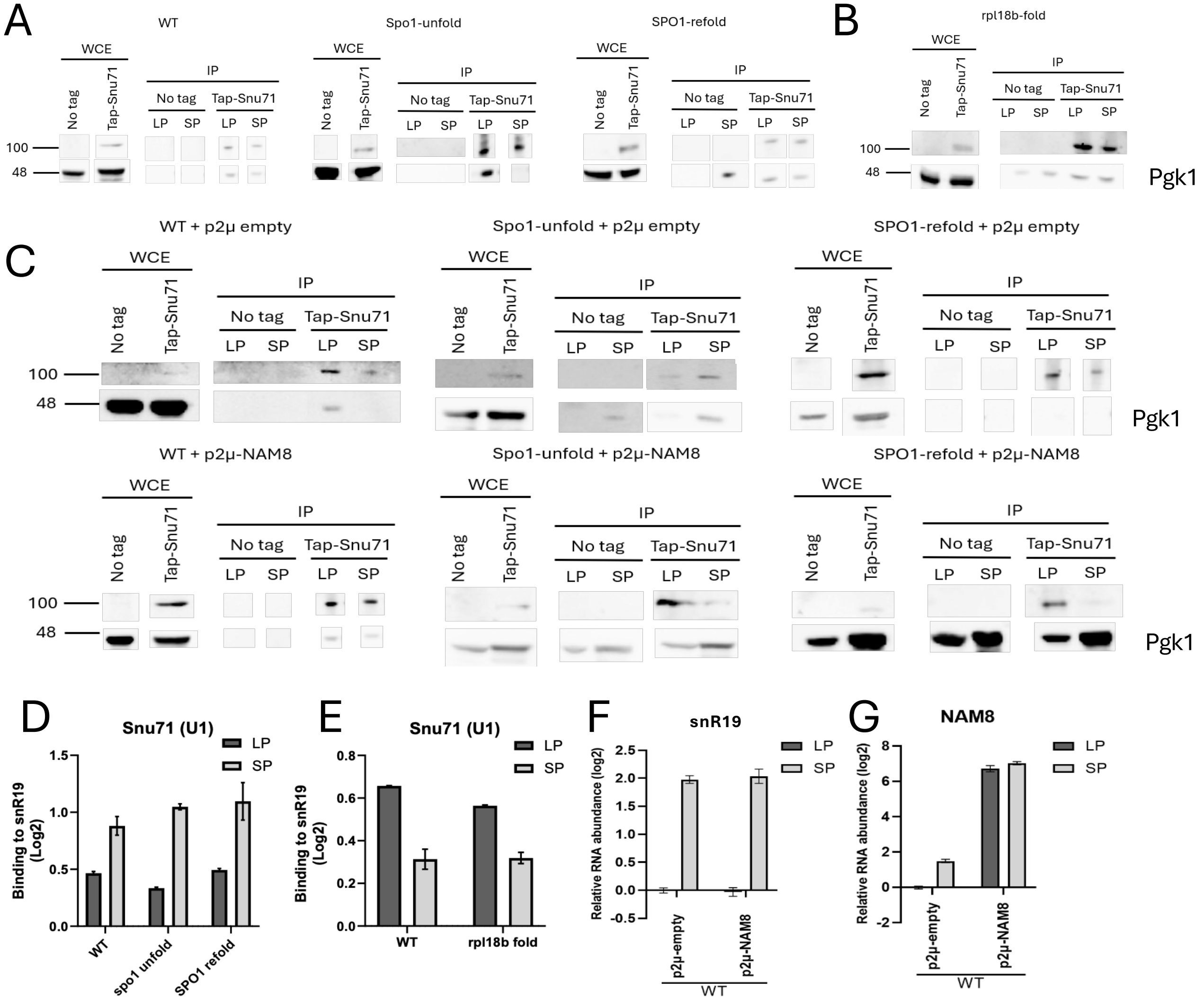


**Figure S4. Validation of U1 snRNP (Snu71) RNA immunoprecipitation and normalization controls, related to Figures 3–5.** (**A–C**) Western blot validation of TAP-Snu71 immunoprecipitation. Untagged control cells and cells expressing TAP-tagged Snu71, a U1 snRNP component, were grown to log phase (LP) or 48 h stationary phase (SP), formaldehyde-crosslinked, and subjected to immunoprecipitation. Whole-cell extract (WCE) and immunoprecipitated (IP) fractions were analyzed by western blot. TAP-Snu71 was enriched in IP fractions relative to untagged controls in SPO1 gate-mutant strains (WT, spo1-unfold, and SPO1-refold) (**A**), the rpl18b-fold strain (**B**), and SPO1 gate-mutant strains carrying empty vector or a *NAM8* overexpression plasmid (**C**). Pgk1 serves as a WCE loading control. Molecular weight markers are indicated in kDa. (**D–E**) snR19 recovery controls for U1 snRNP immunoprecipitation. The amount of snR19, the U1 snRNA, co-immunoprecipitated with Snu71 was quantified by RT-qPCR in WT, spo1-unfold, and SPO1-refold strains (**D**), and WT RPL18B and rpl18b-fold strains (**E**). (**F**) snR19 level in *NAM8* overexpression cells. The amount of snR19 RNA was quantified by RT-qPCR in WT cells carrying empty vector or a *NAM8* overexpression plasmid. (**G**) *NAM8* overexpression. The amount of NAM8 RNA was quantified by RT-qPCR in WT cells carrying empty vector or a *NAM8* overexpression plasmid. (**D–G**) RNA and co-immunoprecipitated RNA was quantified by RT-qPCR, normalized to the unaffected noncoding RNAs RPR1 and NME1, and corrected by subtracting signal from untagged controls for the co-immunoprecipitated RNA. Bars show corrected, normalized RNA as mean ± SD from at least three biological replicates. LP, log phase; SP, stationary phase.

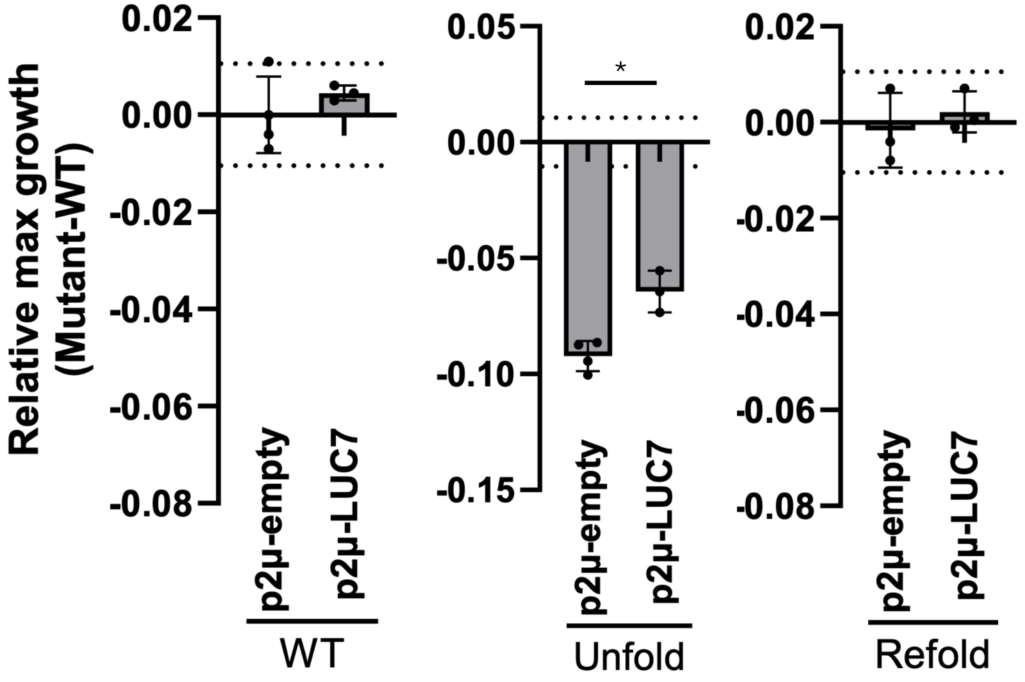


**Figure S5. The general U1 factor Luc7 does not rescue SPO1 gate disruption during starvation growth, related to Figure 5A.** *LUC7* overexpression does not restore the low-dextrose starvation growth defect caused by disruption of the SPO1 5′UTR–intron gate. Relative maximum growth is shown as each strain minus the WT strain carrying the empty vector for WT, spo1-unfold, and SPO1-refold strains carrying empty vector or a *LUC7* overexpression plasmid in amino acid-depleted minimal medium containing 0.1% dextrose. Dotted lines indicate the SD range of WT controls. Bars show mean ± SD from at least three biological replicates. Significance was determined using a two-sided t-test assuming unequal variances; one asterisk, $p < 0.05$.

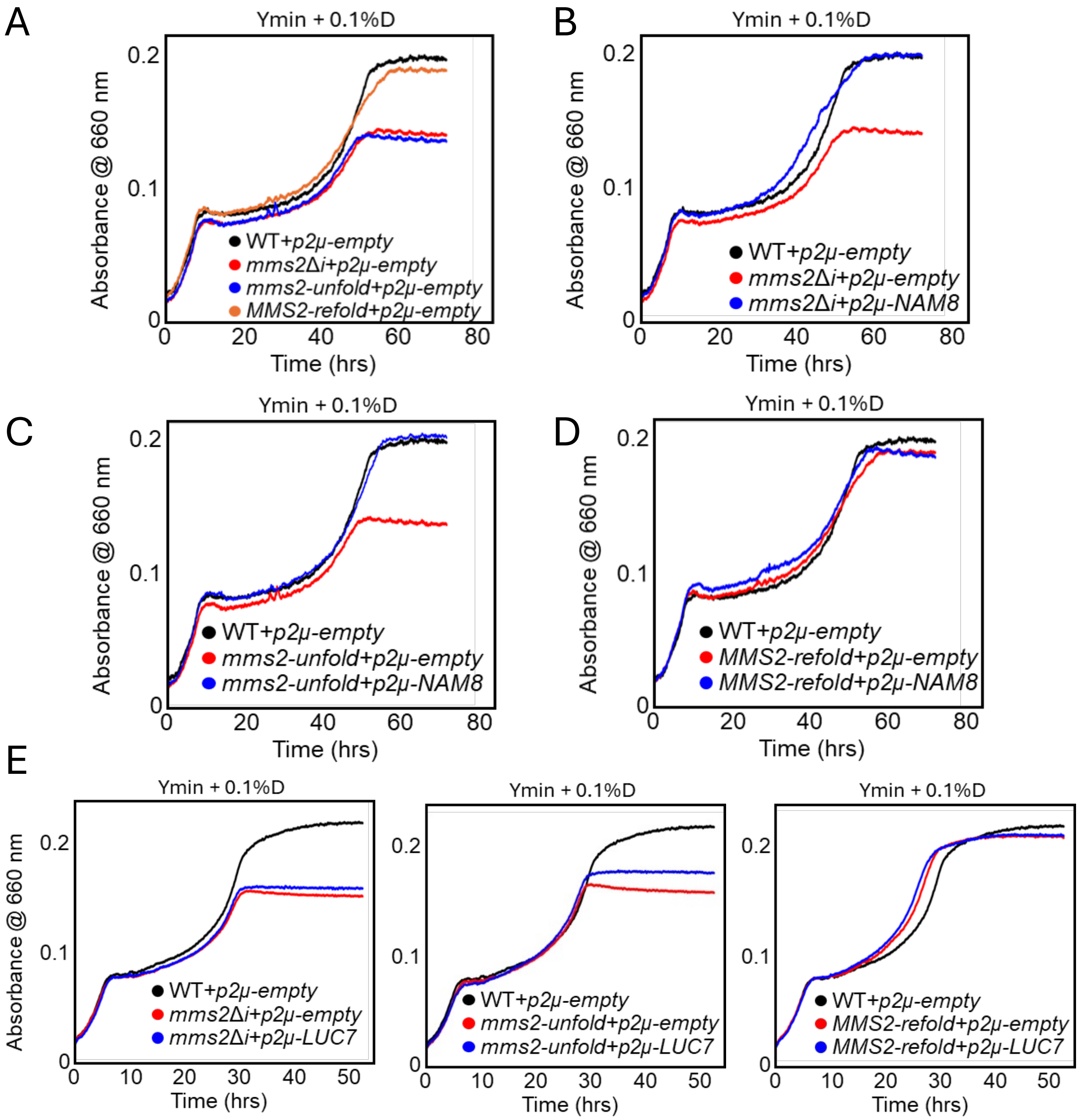


**Figure S6. Nam8, but not the general U1 factor Luc7, restores starvation growth of a second gate-disrupted UpSI, MMS2, related to Figure 5.** (**A**) Growth of MMS2 gate-mutant strains carrying empty vector. WT, *mms2Δi*, *mms2-unfold*, and *MMS2-refold* strains carrying empty vector were grown in amino acid-depleted minimal medium containing 0.1% dextrose. Growth was monitored by absorbance at 660 nm over 72 h. Disruption of the MMS2 5′UTR–intron gate phenocopied intron deletion under low-dextrose starvation conditions, whereas restoring base pairing rescued growth toward the WT trajectory. (**B–D**) *NAM8* overexpression restores starvation growth of *MMS2* intron-deletion and gate-disrupted strains. Growth of *mms2Δi* (**B**), *mms2-unfold* (**C**), and *MMS2-refold* (**D**) strains carrying empty vector or a *NAM8* overexpression plasmid was compared with WT carrying empty vector in amino acid-depleted minimal medium containing 0.1% dextrose. (**E**) *LUC7* overexpression does not restore starvation growth of *MMS2* intron-deletion or gate-disrupted strains. Growth of *mms2Δi*, *mms2-unfold*, and *MMS2-refold* strains carrying empty vector or a *LUC7* overexpression plasmid was compared with WT carrying empty vector in amino acid-depleted minimal medium containing 0.1% dextrose. (**A–E**) Curves show mean OD660 from at least three biological replicates.
